# TRITEX: chromosome-scale sequence assembly of Triticeae genomes with open-source tools

**DOI:** 10.1101/631648

**Authors:** Cécile Monat, Sudharsan Padmarasu, Thomas Lux, Thomas Wicker, Heidrun Gundlach, Axel Himmelbach, Jennifer Ens, Chengdao Li, Gary J. Muehlbauer, Alan H. Schulman, Robbie Waugh, Ilka Braumann, Curtis Pozniak, Uwe Scholz, Klaus F. X. Mayer, Manuel Spannagl, Nils Stein, Martin Mascher

**Affiliations:** Leibniz Institute of Plant Genetics and Crop Plant Research (IPK) Gatersleben, Seeland, Germany; PGSB – Plant Genome and Systems Biology, Helmholtz Center Munich – German Research Center for Environmental Health, Neuherberg, Germany; Department of Plant and Microbial Biology, University of Zurich, Zurich, Switzerland; Department of Plant Sciences, University of Saskatchewan, Saskatoon, Canada; Western Barley Genetics Alliance, School of Veterinary and Life Sciences (VLS), Murdoch University, Murdoch, WA, Australia; Hubei Collaborative Innovation Center for Grain Industry / School of Agriculture, Yangtze University, Jingzhou, China; Department of Agronomy and Plant Genetics & Department of Plant and Microbial Biology, University of Minnesota, St. Paul, Minnesota, USA; Green Technology, Natural Resources Institute (Luke), Viikki Plant Science Centre, and Institute of Biotechnology, University of Helsinki, Helsinki, Finland; The James Hutton Institute, Dundee, UK; School of Life Sciences, University of Dundee, Dundee, UK; Carlsberg Research Laboratory, Copenhagen, Denmark; School of Life Sciences Weihenstephan, Technical University of Munich, Germany; Department of Crop Sciences, Center for Integrated Breeding Research (CiBreed), Georg-August-University Göttingen, Göttingen, Germany; German Centre for Integrative Biodiversity Research (iDiv) Halle-Jena-Leipzig, Leipzig, Germany

**Author notes:** Correspondence should be addressed to Martin Mascher or Nils Stein.

## Abstract

Chromosome-scale genome sequence assemblies underpin pan-genomic studies. Recent genome assembly efforts in the large-genome Triticeae crops wheat and barley have relied on the commercial closed-source assembly algorithm DeNovoMagic. We have developed TRITEX, an open-source computational workflow that combines paired-end, mate-pair, 10X Genomics linked-read with chromosome conformation capture sequencing data to construct sequence scaffolds with megabase-scale contiguity ordered into chromosomal pseudomolecules. We evaluated the performance of TRITEX on publicly available sequence data of tetraploid wild emmer and hexaploid bread wheat, and constructed an improved annotated reference genome sequence assembly of the barley cultivar Morex as a community resource.

## Introduction

The Triticeae species wheat and barley were among the founder crops of Neolithic agriculture in Western Asia and continue to dominate agriculture in temperate regions of the world to the present day. Large genome sizes, high content of transposable elements (TEs) and polyploidy (in the case of wheat) have long impeded genome assembly projects in the Triticeae [1, 2]. Recently, chromosome-scale reference sequence assemblies have come available for barley (*Hordeum vulgare*) [3], hexaploid bread wheat (*Triticum aestivum*) [4], tetraploid durum wheat (*T. turgidum ssp. durum*) [5] as well as the wheat wild relatives *Aegilops tauschii* (wheat D genome progenitor) [6], T. urartu (wheat A genome progenitor) [7] and *T. turgidum* ssp. *dicoccoides* (wild emmer wheat, AB genome) [8]. The genome projects of barley, bread wheat and the A and D genome progenitors had initially followed the hierarchical shotgun approach as had been employed by the human genome project [9], but adopted second-generation sequencing methods for sequencing as they became available [10]. Assembling bacterial artificial chromosomes (BACs) guided by a physical map yielded megabase-sized scaffolds [3, 11], which were then arranged into chromosomal super-scaffolds (so-called “pseudomolecules”) by long-range linkage information afforded by ultra-dense genetic maps [12, 13], chromosome conformation capture sequencing (Hi-C) [14, 15] or Bionano optical mapping [16]. However, BAC-by-BAC assembly is laborious and time-consuming [3], and has become an obsolete method of sequence assembly.

The wild emmer wheat, and subsequently the bread and durum wheat, genome projects [4, 5, 17] used a whole-genome shotgun (WGS) approach based on Illumina short-read sequencing of shotgun libraries with multiple insert sizes. Within months, a fully annotated, highly contiguous sequences was assembly, capturing the full organizational context of the 21 wheat chromosomes, some of which have been validated using other approaches [18]. Despite being robust, the assembly algorithm used in these projects was closed-source [19], potentially limiting its application to the broader community. Indeed, efforts to develop a low-cost, open source alternative are still required to allow assembly of multiple genomes within a species to comparable contiguity. Short-read assemblies of the wheat genome have been generated by open-source alternatives such as w2rap [20] or Meraculous [13]. In addition, long-read assemblies have been generated for *Ae. tauschii* [21] and bread wheat [22]. But still the contiguity of these assemblies is lower than that of the scaffolds constructed using the DeNovoMagic algorithm. Another important concern is the high computational cost for a long-read (hybrid) assembly, estimated at 470,000 CPU hours or 6.5 months in wall-clock time [22].

We have recently outlined a proposal for pan-genomics in barley [23]. A cornerstone of our strategy is the construction of high-quality sequence assemblies for multiple genotypes representative of major germplasm groups. Similar projects are under way in bread wheat (http://www.10wheatgenomes.com). An open-source assembly pipeline with comparable accuracy, completeness and speed similar to available commercial platforms would greatly reduce the cost per assembled genome, thus extending the scope of pan-genome projects in the Triticeae.

Here, we report on the development of a computational pipeline for chromosome-scale sequence assembly of wheat and barley genomes. We evaluate the performance of the pipeline (which we named TRITEX) by re-assembling the raw data used for the wild emmer [8] and bread wheat [24] reference genome assemblies and compare our assemblies to the those constructed with a commercial platform. Furthermore, we used TRITEX to generate an improved annotated reference genome assembly for barley cv. Morex as an important resource for the barley research community.

## Results

### Overview of the workflow

We begin with a description of our workflow, its input datasets (**Table 1)** and a description of the expected outcome of each component (**Table 2**). For the sake of exposition, we illustrate our method by presenting results for wheat and barley, which will be described in greater detail below.

**Table 1:** Input datasets for TRITEX.

| Name | Library type (number<sup>1</sup>) | Insert size | Read length | Coverage<sup>2</sup> |
| --- | --- | --- | --- | --- |
| PE450 | PCR-free paired-end (2) | 400-470 bp | 2x250 bp | 70x |
| PE800 | PCR-free paired-end (2) | 700-800 bp | 2x150 bp | 30x |
| MP3 | Nextera mate-pair (2) | 2-4 kb | 2x150 bp | 30x |
| MP6 | Nextera mate-pair (2) | 5-7 kb | 2x150 bp | 30x |
| MP9 | Nextera mate-pair (2) | 8-10 kb | 2x150 bp | 30x |
| 10X | 10X Chromium (2) |  | 2x150 bp | 30x |
| Hi-C | TCC [90] or in-situ Hi-C [91] (1) |  | 2x100 bp | 200 – 400 million read pairs |

<sup>1</sup>Number of independent libraries to be prepared.
<sup>2</sup>Haploid genome coverage for paired-end, mate-pair and 10X libraries. As Hi-C analysis is count-based, read numbers are more relevant than sequence amount.

**Table 2:** Overview of the TRITEX pipeline.

|  | **Step<sup>1</sup>** | **Software** | **Input** | **Output** |
| --- | --- | --- | --- | --- |
| **1** | Read merging | BBMerge [26] | PE450 read pairs | Merged PE450 reads |
| **2** | PE450 error-correction | BFC [27] | Merged PE450 reads | Corrected PE450 reads<br>Hash table of k-mer counts |
| **3.1** | Unitig assembly | Minia3 [29, 30] | Corrected PE450 reads | Unitigs |
| **3.2** | Error-correction of<br>PE800 and MP reads | BFC [27]<br>Cutadapt [60],<br>NxTrim [61] | PE800, MP3, MP6, MP9 reads<br>Hash table of k-mer count (step 2) | Corrected PE800, MP3, MP6 and MP9 reads |
| **4** | Scaffolding | SOAPDenovo2 [62] | Unitigs<br>Corrected PE800, MP3, MP6, MP9 reads | Scaffolds |
| **5** | Gap-filling | Gapcloser [62] | Scaffolds<br>Corrected PE450 reads | Scaffolds after gap-filling |
| **6.1** | Alignment of 10X reads | Minimap2 [36],<br>cutadapt [60],<br>SAMtools [63],<br>BEDtools [64],<br>custom scripts | Scaffolds after gap-filling<br>10X reads | 10X alignment records |
| **6.2** | Alignment of Hi-C reads | As in 6.2<br>EMBOSS [66] | Scaffolds after gap-filling<br>Hi-C reads | Hi-C alignment records |
| **6.3** | Alignment of genetic<br>markers | Minimap2 [36] | Scaffolds after gap-filling<br>Marker sequences | Marker alignment records |
| **7** | Pseudomolecule<br>construction | Custom R scripts | Scaffolds after gap-filling<br>10X alignment records<br>Hi-C alignment records<br>Marker alignment records | Pseudomolecules<br>Hi-C contact maps |

<sup>1</sup>Steps with identical leading digits can be run in parallel.

Our pipeline uses the same input datasets generated for DeNovoMagic assemblies reported by Avni et al. [17] and the International Wheat Genome Sequencing Consortium (IWGSC) [4]. The key parameters are two types of paired-end libraries (PE450 and PE800), three types of mate-pair libraries (size ranges: 2-4 kb [MP3], 5-7 kb [MP6] and 8-10 kb [MP9]), 10X Chromium libraries, and Hi-C data as listed in **Table 1**. We show below that certain library types can be omitted in our approach without greatly compromising assembly contiguity.

A critical component of the “sequencing recipe” are PCR-free Illumina shotgun libraries with a tight insert size distribution in the range of 400-500 bp and sequenced with 250 bp paired-end reads. These were merged with standard tools such as PEAR [25] or BBMerge [26] to yield long single-end reads with a mean fragment size of ~450 bp and are subsequently error-corrected with BFC [27]. These elongated short reads allow the use of longer k-mers (i.e. short sequence fragments of fixed length) during the assembly process. Estimates of expected assembly size based on k-mer cardinalities [28] support the notion that longer k-mers achieve much better genome representation in wheat and barley (Fig. 1a). The k-mer size for many assemblers is limited. For example, the maximum k-mer size of SOAPDenovo2 is limited to 127 bp. We thus selected Minia3 [29, 30], an assembler capable of using k-mers of arbitrary size.

**Figure 1:**
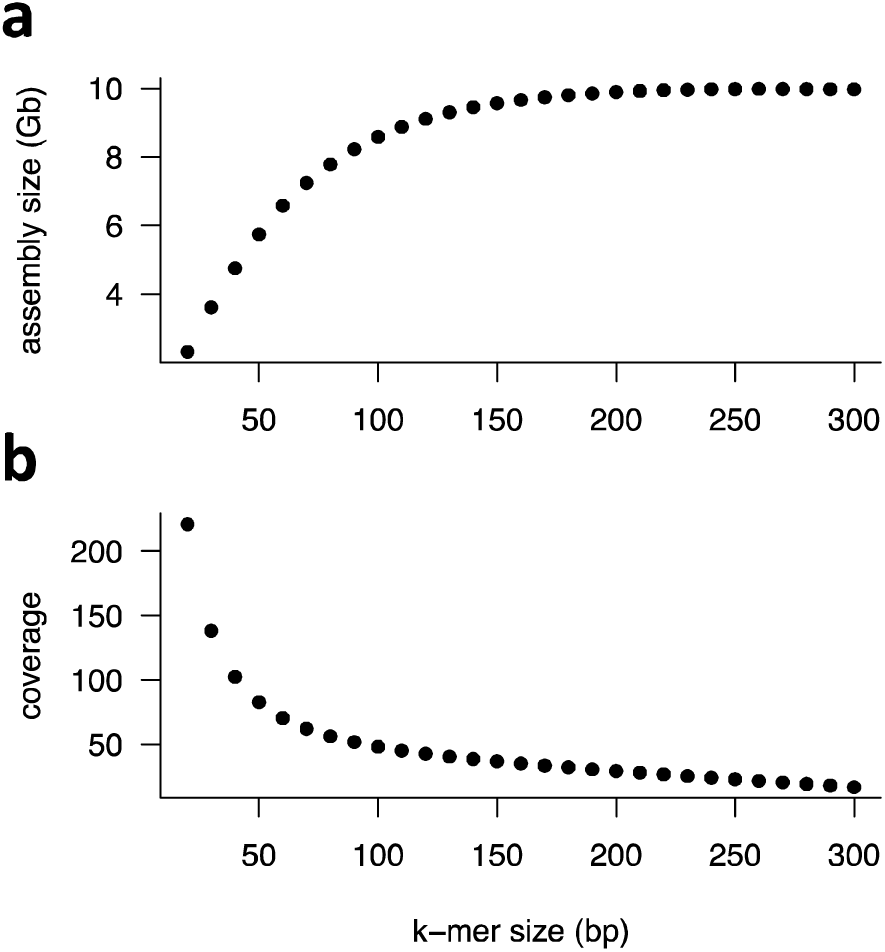
Estimate of assembly size and k-mer coverage as a function of k-mer size. Assembly size **(a)** and k-mer coverage **(b)** were estimated from error-corrected PE450 used for Zavitan unitig assembly based on k-mer cardinalities using NtCard [92] and Kmerstream [28].

However, one disadvantage of using large k-mer sizes is the lower genome coverage (**Fig. 1b**) as a consequence of sequencing errors, resulting in random coverage gaps. To overcome this drawback, we adopted the iterative multi k-mer approach of the GATB-Minia pipeline https://github.com/GATB/gatb-minia-pipeline). In the initial iteration, an assembly at k-mer size 100 is made from the error-corrected PE450 reads. Subsequent iterations take as input the PE450 reads and assembly constructed in the previous iteration. This procedure is repeated for k-mer sizes 200, 300, 350, 400, 450 and 500. The unitigs of the final iteration achieve an N50 of about 20 – 30 kb (**Table 3**). A single iteration takes about one day for barley and three days for hexaploid wheat.

**Table 3:** Assembly statistics for Zavitan and Chinese Spring.

|  | **Zavitan** |  | **Chinese Spring** |  |
| --- | --- | --- | --- | --- |
|  | TRITEX | Avni et al. [8] | TRITEX | IWGSC [4] |
| **Unitig assembly size** | 10.8 Gb |  | 15.1 Gb |  |
| **Unitig N50** | 21.7 kb |  | 21.4 kb |  |
| **Unitig N90** | 1.5 kb |  | 1.7 kb |  |
| **Assembled sequence in contigs >= 1 kb** | 10.0 Gb |  | 14.0 Gb |  |
| **Assembled sequence in contigs >= 10 kb** | 7.8 Gb |  | 10.8 Gb |  |
| **Scaffold assembly size** | 11.1 Gb | 10.5 Gb | 15.7 Gb | 14.5 Gb |
| **Scaffold N50** | 1.3 Mb | 7.0 Mb | 2.3 Mb | 7.0 Mb |
| **Scaffold N90** | 97 kb | 1.2 Mb | 281 kb | 1.2 Mb |
| **Assembled sequence in scaffolds >= 1 kb** | 10.4 Gb | 10.5 Gb | 14.8 Gb | 14.5 Gb |
| **Assembled sequence in scaffolds >= 1 Mb** | 6.7 Gb | 9.6 Gb | 11.9 Gb | 13.4 Gb |
| **Unfilled internal gaps** | 209 Mb (1.9 %) | 171 Mb (1.6 %) | 476 Mb (3.0 %) | 262 Mb (1.8 %) |

The unitigs of the final iteration are used as input for scaffolding with the PE800, MP3, MP6 and MP9 libraries using SOAPDenovo2 [6]. This yields assemblies with an N50 beyond 1 Mb (**Table 3, 4**). After gap-filling with GapCloser, about 1 – 5 % internal gaps in scaffolds remain (**Tables 3, 4**). Alignments of 10X and Hi-C reads and genetic markers to the scaffolds are imported into R [31] and custom scripts were developed to identify and correct mis-assemblies, to construct super-scaffolds and to build pseudomolecules. Both super-scaffolding with 10X data and pseudomolecule construction use the POPSEQ genetic maps of barley [12] and wheat [13] to guide the assignment of scaffolds to chromosomes and to discard spurious links between unlinked regions. We note that the omission of the PE800, MP3 and MP6 libraries (i.e. using only the MP9 library for mate-pair scaffolding) resulted in assemblies of comparable contiguity and genome representation in barley (**Table 4**). If this slightly reduced contiguity is acceptable for downstream application, the cost for data generation can be reduced by about 20%.

**Table 4:**
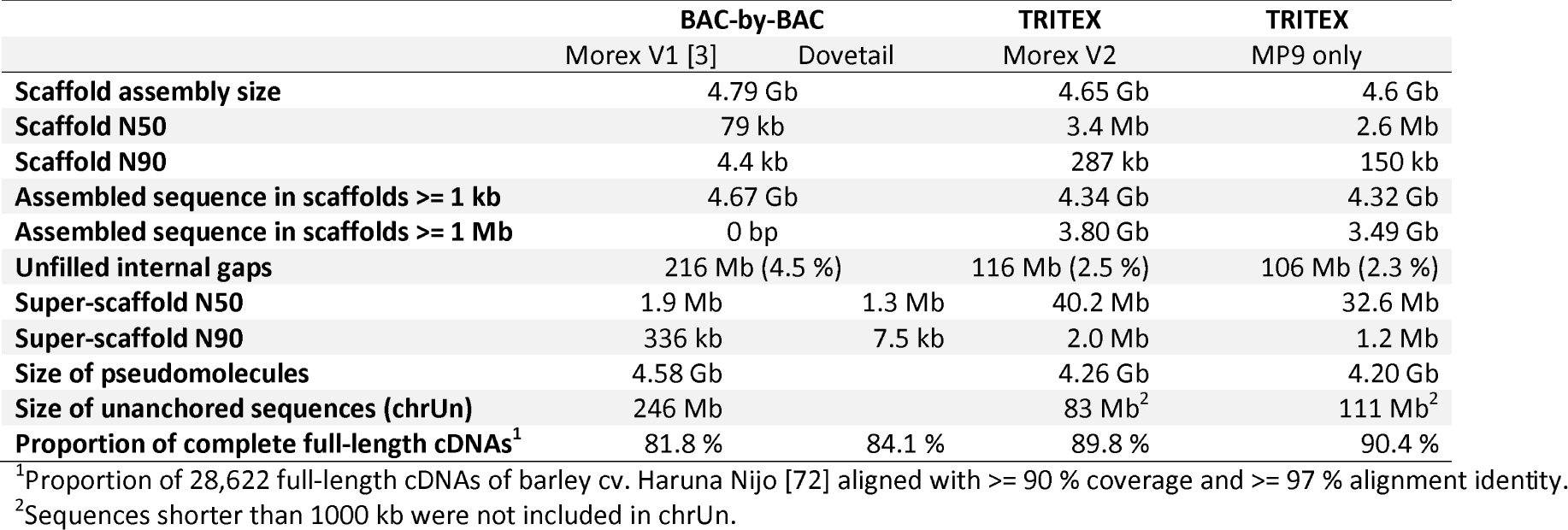
Comparison of different assemblies of barley cv. Morex.

Scaffolding can introduce false joins between unlinked sequences [32] that need to be broken to construct correct chromosomal pseudomolecules [33]. Physical coverage with 10X reads is used to detect and correct mis-joins introduced during either unitig construction or scaffolding (**Fig. 2**). The corrected scaffolds are used as input for super-scaffolding with 10X data using a custom graph-based method (see Methods section for details). These super-scaffolds are then ordered and oriented along the chromosomes using Hi-C data using the method of Beier et al. [34]. Once scaffolds have been arranged into chromosomal pseudomolecules, contact matrices for each chromosome are plotted as heat maps. Visual inspection of these matrices can reveal further assembly errors such as remaining chimeras or misoriented (blocks of) super-scaffolds (**Fig. 3**). After correction of these errors, the Hi-C maps are updated and cycles of assembly-inspection-correction are repeated until all mis-assemblies have been eliminated and contact matrices show the expected Rabl configuration [3] (strong main diagonal / weak anti-diagonal). We found that pseudomolecules constructed from corrected super-scaffolds contain in the range of 10 to 20 mis-assemblies, which were eliminated in a single correction cycle. Without 10X data, i.e. using only Hi-C data for spotting misassemblies as in the case of published wheat reference genomes [4, 17], more curation cycles were required.

**Figure 2:**
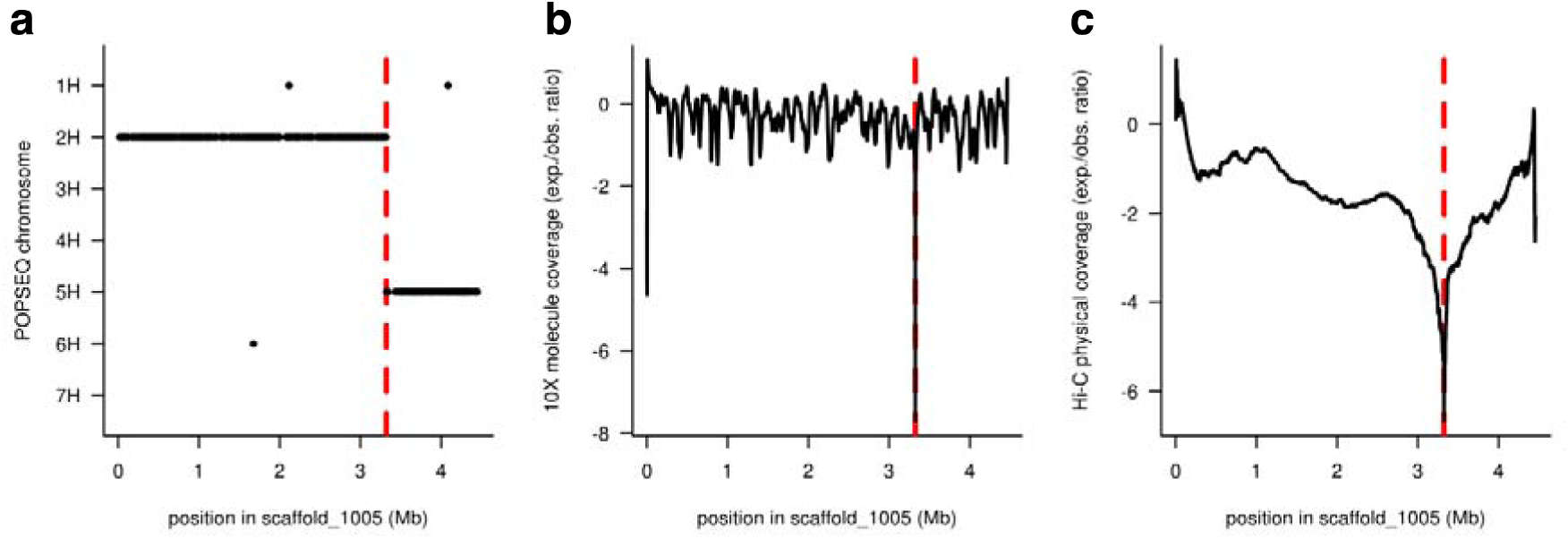
Example of a chimeric scaffold. The chimeric nature of a sequence scaffold joining two unlinked sequences originating from barley chromosomes 2H and 5H is supported by multiple lines of evidence. **(a)** Genetic chromosome assignments of marker sequences aligned to scaffold_1005. **(b)** 10× molecule coverage. **(c)** Physical Hi-C coverage. Coverage in panels **(b)** and **(c)** was normalized for distance from the scaffold ends and the log2-fold observed vs. expected ratio was plotted. The red, dotted lines mark the breakpoint at 3.32 Mb.

**Figure 3:**
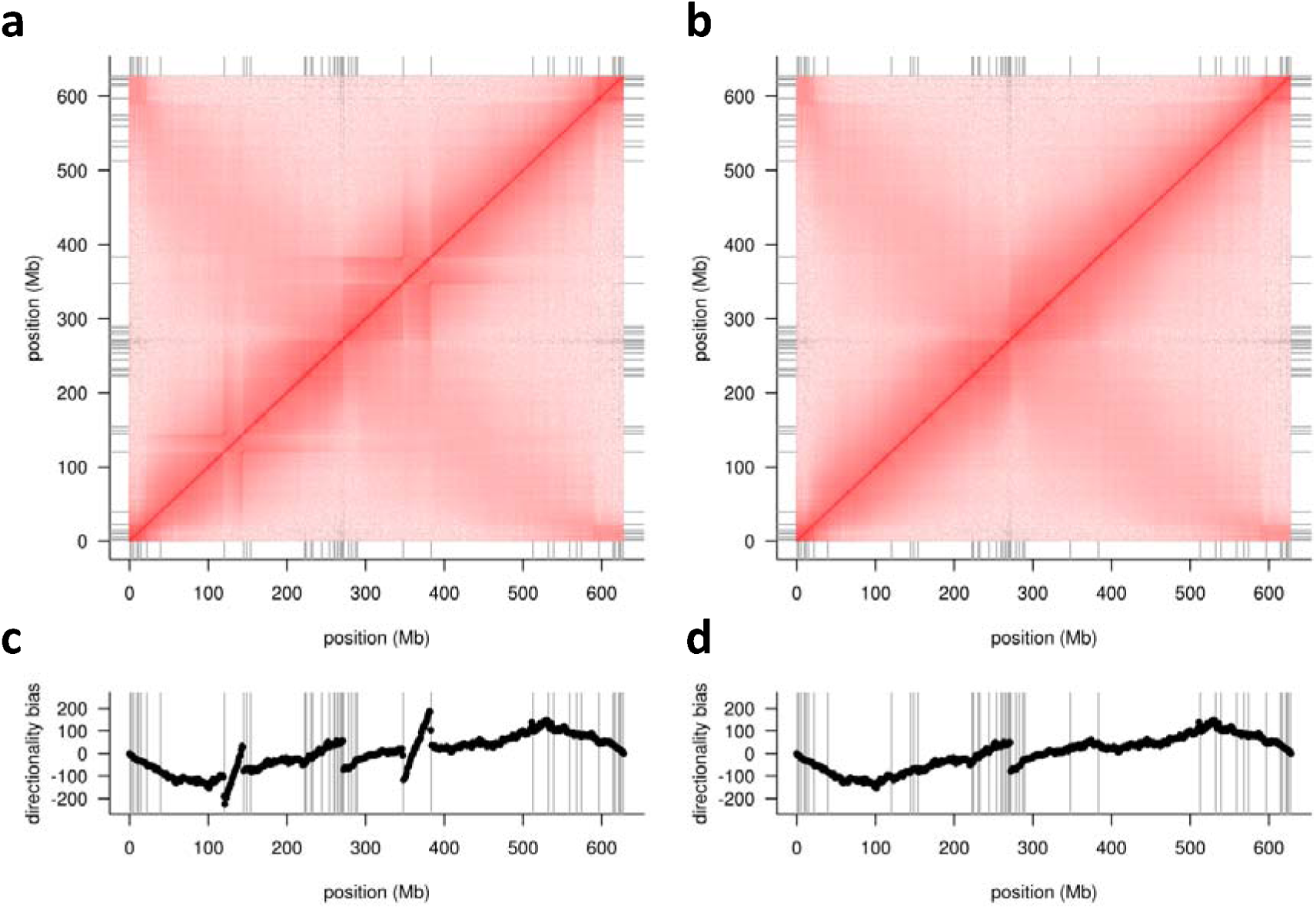
Example of errors in scaffold orientation. The top panels show the Hi-C contact matrix for barley chromosome 3H before **(a)** and after **(b)** manual correction. The bottom panels show the directionality biases in the Hi-C data as defined by Himmelbach et al. [93] before **(c)** and after **(d)** manual correction. Two inverted scaffolds are evident as deviations from the expected Rabl configuration [3] and as diagonals bounded by discontinuities in the directionality biases.

Assuming all input datasets (**Table 1**) are in place, the entire TRITEX workflow can be completed in three to four weeks for barley and four to six weeks for hexaploid wheat, allowing for some delays in the completion of hands-on steps (mainly inspection of intermediate results and curation of pseudomolecules). We believe that despite our detailed user guide (available at https://tritexassembly.bitbucket.io), completing a TRITEX assembly would be a rather arduous task for a scientist inexperienced in either plant genome assembly or practical bioinformatics, unless guided by an expert in plant genomics. A UNIX server with at least 1 TB of main memory is needed to complete scaffold construction for bread wheat. Much wall-clock time (about 1 week for barley [5 Gb genome) and about 3 weeks for bread wheat [16 Gb genome]) is spent for unitig assembly with Minia3. Fortunately, the main memory consumption of Minia3 is low (50 GB). Thus, assemblies of multiple genotypes (a typical use case in a pan-genome project) can be run in parallel.

### Re-assembly of wild emmer and bread wheat and comparison to published assemblies

We downloaded the paired-end and mate-pair reads used for the DeNovoMagic assemblies of wild emmer wheat accession Zavitan [17] and bread wheat cultivar Chinese Spring [4] (referred to as the IWGSC whole-genome assembly in Table S2 of [4]) and ran TRITEX until the gap-filling step (step 2 in **Table 2**). The metrics of TRITEX assemblies were inferior to those of DeNovoMagic (**Table 3**). Still, the contiguity of the TRITEX assemblies was in the megabase range, and it was clearly superior to the BAC-by-BAC assembly of a single wheat chromosome (3B [11], N50: 892 kb). Visual inspection of alignments between the TRITEX and DeNovoMagic assemblies indicated a high concordance between them (**Fig. 4**). To assess the accuracy of the TRITEX assemblies at a genome-wide scale, we compared the TRITEX scaffolds to published assemblies produced by the DeNovoMagic algorithm. These assemblies had been independently validated by complementary sequence and mapping resources [4, 17, 35]. We divided the TRITEX scaffolds into non-overlapping 10 kb fragments, aligned them to the DeNovoMagic assemblies with Minimap2 [36] and measured the collinearity of the alignments. The average Pearson correlation of fragment positions in the TRITEX scaffold and their aligned positions in the DeNovoMagic scaffolds was 0.998 for Chinese Spring and 0.999 for Zavitan. Across all Chinese Spring (Zavitan) scaffold pairs with at least 100 kb of aligned sequences, 99.96 % (99.99 %) of aligned fragment sequences were mapped in the same orientation. These results support a very high concordance in the local order and orientation of sequences between the TRITEX and previous assembly efforts.

**Figure 4:**
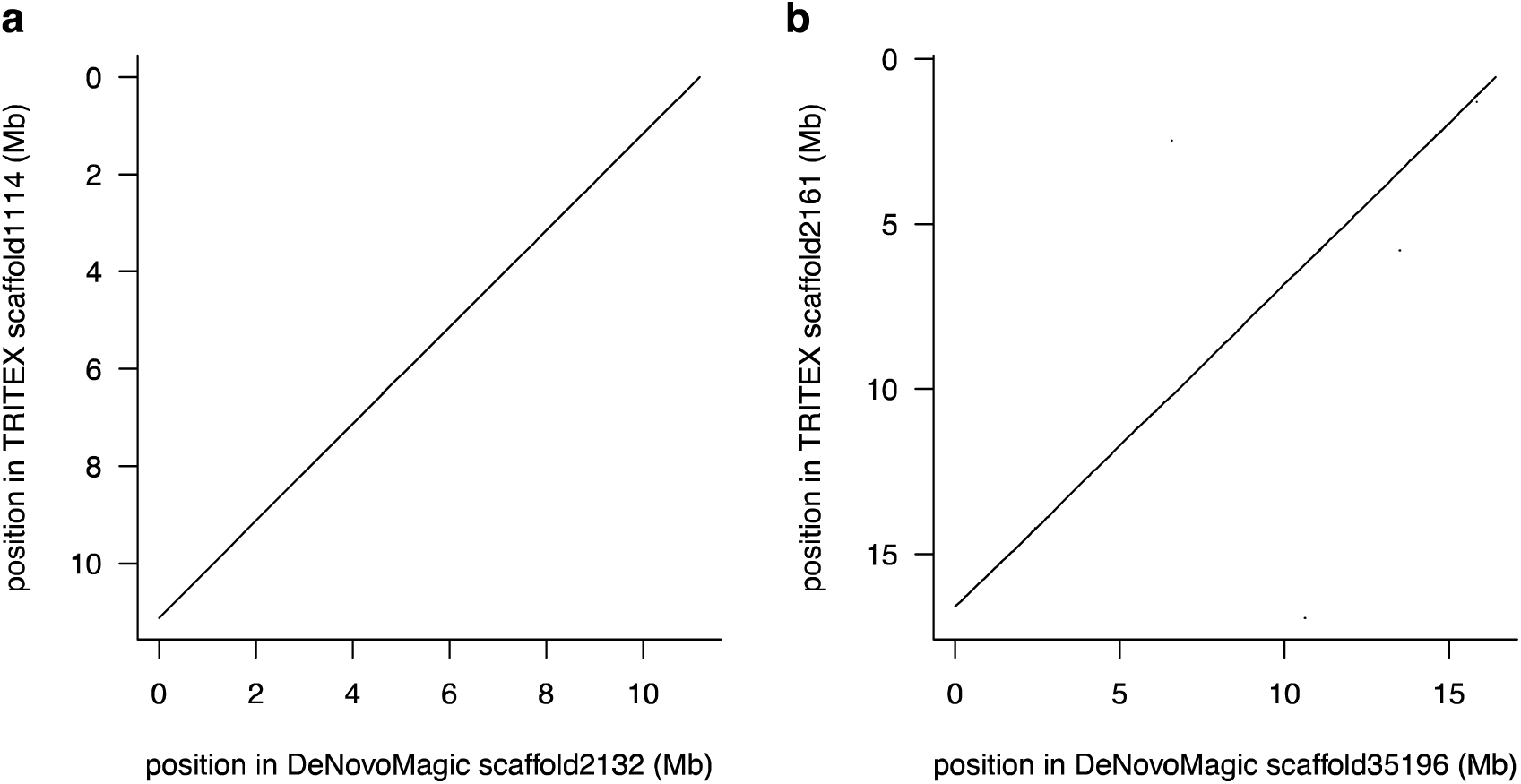
Collinearity between TRITEX and DeNovoMagic assemblies of wheat. Dotplots showing the longest alignments between scaffold pairs of the TRITEX and DeNovoMagic assemblies of Zavitan **(a)** and Chinese Spring **(b)**, respectively. Alignments were done with Minimap2 [36].

To assess the completeness of the TRITEX assembly of Chinese Spring, we determined the representation of two transcript resources: the IWGSC gene models [4] and the full-length cDNAs of Mochida et al. [37]. The proportion of completely represented transcripts in the TRITEX assembly was very similar to the IWGSC RefSeq and substantially higher than in the w2rap assembly [20] and the PacBio hybrid assembly of Zimin et al. [22] (Table 5). Note that the IWGSC RefSeq gene models are likely to have a bias for the TRITEX assembly, which was generated from the same input data, but it is not evident how the Sanger-sequenced full-length cDNAs might favor a certain assembly.

**Table 5:** Chinese Spring transcript alignment statistics.

| Transcript dataset | No. of transcripts | Assembly | Proportion of complete transcripts<sup>1</sup> |
| --- | --- | --- | --- |
| IWGSC v1.0 transcripts [4] | 269,583 | TRITEX | 96.2 % |
|  |  | IWGSC [4] | 97.0 % |
|  |  | Clavijo et al. [20] | 87.8 % |
|  |  | Zimin et al. [22] | 88.5 % |
| Full-length cDNAs [37] | 6,137 | TRITEX | 97.1 % |
|  |  | IWGSC [4] | 96.3 % |
|  |  | Clavijo et al. [20] | 91.6 % |
|  |  | Zimin et al. [22] | 85.4 % |

<sup>1</sup> Proportion of transcripts with at least one alignment with $\geq 99$ % coverage and $\geq 99$ % identity (for IWGSC transcripts) or with $\geq 90$ % coverage and $\geq 99$ % identity (for full-length cDNAs). Alignments were done with GMAP [70].

A comparison of the TRITEX assemblies of Zavitan and Chinese Spring in comparison to their published counterparts revealed a higher proportion of sequence gaps (Table 3). We speculate that sequence gaps may arise because highly similar copies of transposable elements (TEs) cannot be resolved. To test this hypothesis, we analyzed the representation of two TE families, RLC_Angela and RLC_Sabrina [38], in the TRITEX and IWGSC WGA assemblies of Chinese Spring. Despite having similar assembly sizes (Table 3), we identified substantially fewer full-length RLC_Angela elements in the TRITEX assembly, whereas the numbers of RLC_Sabrina elements matched closely in both assemblies (Table S1). RLC_Angela is considered a recently active (i.e. transposing) family, whereas RLC_Sabrina has been inactive for a long time (Thomas Wicker, unpublished results). Consistent with the expectation that younger elements, which inserted recently and have highly similar copies elsewhere in the genomes, are not well assembled, the age distribution of RLC_Angela is skewed for older elements. By contrast, no such bias is seen for RLC_Sabrina (Figure S1). In summary, the TRITEX assembly of Chinese Spring has fewer complete TEs, indicating that the DenovoMagic algorithm may make better use of mate-pair or PE450 data to close gaps in TEs.

### An improved barley reference genome assembly

Prompted by the encouraging assembly results for wheat, we decided to employ the TRITEX pipeline to construct a second version reference genome assembly of barley cv. Morex. The need for an improved assembly arose from shortcomings of the BAC-based reference sequence [3] including (1) large sequence gaps, (2) redundancies and (3) local mis-assemblies.

First, gaps in the physical map or failed BAC assemblies result in gaps in the assembled sequence that may contain important genes. During the process of pseudomolecule construction [34], we attempted to “rescue” missed genes by adding sequences from a WGS draft assembly of Morex [39]. However, the short WGS contigs do not provide the local sequence context of genes and may not even contain full-length gene sequences. Second, it was necessary to merge sequences from individually assembled BAC clones [10]. during pseudomolecule construction [34]. Megabase-sized sequence scaffolds representing physical contigs of BACs were constructed using a complex, multi-tiered method that employed heuristic approaches to distinguish true sequence overlaps from alignments caused by highly similar copies of transposable elements [34]. Nevertheless, self-alignment of the pseudomolecule at a high identity threshold (minimum alignment length: 5 kb, minimum alignment identity: 99.5 %) resulted in a substantial proportion (4.4 %) of undetected overlaps between adjacent BACs. Similar results were obtained for the first version of the BAC-based maize reference genome [40] (1.2 %) and the 3B pseudomolecule of Choulet et al. [11] (7.4 %) using the same alignment thresholds.

Third, individual BAC clones were rarely represented by a single sequence scaffold even after scaffolding with mate-pair data [10]. At the time barley BAC sequencing was performed, methods with a sufficient density and resolution to order and orient sequence scaffolds within 100 kb were not available. Hence, our solution [34] was to place sequence scaffolds originating from the BAC clone in arbitrary order and orientation into the pseudomolecule, thus introducing many local assembly errors at the sub-BAC scale.

Our results in wheat led us to expect that a TRITEX assembly of the Morex genome would overcome the limitations inherent to the BAC-by-BAC approach. To construct a TRITEX assembly of the Morex genome, we obtained the datasets as detailed in **Table 1**. New paired-end, mate-pair and 10X libraries were constructed and sequenced. The Hi-C data of Mascher et al. [3] were used for pseudomolecule construction. The assembly metrics of the Morex TRITEX assembly greatly exceeded those of the BAC-by-BAC assembly. Notably, the proportion of completely aligned full-length cDNAs improved by about nine percentage points (**Table 4**) compared to the BAC-by-BAC assembly.

To ascertain the correct local order and orientation of sequence scaffolds, we compared the TRITEX super-scaffolds to three complementary resources: (i) the first version (V1) pseudomolecules, (ii) the BAC-by-BAC assembly improved by super-scaffolding based on newly collected in-vitro proximity ligation and (iii) the genome-wide optical map of Morex. First, visual inspection of alignments confirmed the expected discordances at the sub-BAC level, but showed good collinearity at the megabase-scale (**Fig. 5a**). We used the same approach as for the comparison to the published wheat assemblies by aligning 10 kb fragments. The Pearson correlation between TRITEX fragments and their aligned positions in the Morex V1 pseudomolecules was 0.927, reflecting a breakdown of collinearity at finer resolution. The orientations in the pseudomolecules of fragments originating from the TRITEX super-scaffolds was highly discordant: on average, only 63 % of aligned sequence in TRITEX/Morex V1 scaffold pairs was in concordant orientation.

**Figure 5:**
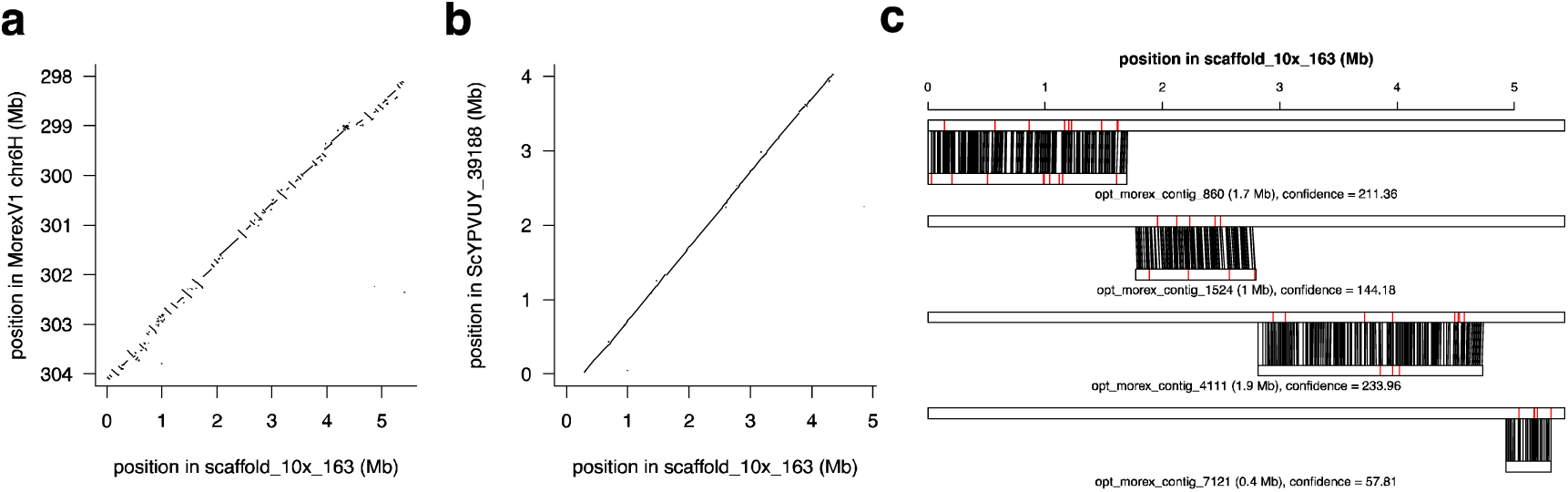
Morex V2 assembly validated by complementary resources. Morex scaffold_10×_163 was aligned to the Morex V1 assembly **(a)**, the Dovetail assembly of Morex **(b)** and the genome-wide optical map of Morex **(c)**. Sequence alignments are shown as dot plots **(a, b)**. Panel **(c)** shows the alignment of four optical contigs to scaffold_10×_163. Single aligned restriction sites are connected by black lines. Red lines indicate unaligned restriction sites in either the sequence scaffold or the optical contig.

Second, we compared the TRITEX super-scaffolds to an improved version of the BAC-by-BAC assemblies of Mascher et al. [3]. Before the development of TRITEX, we had attempted to order and orient BAC sequence scaffolds using the Dovetail method. This involved in-vitro proximity ligation sequencing (Chicago) followed by scaffolding with the HiRise assembler [41]. Visual inspection of alignments between TRITEX super-scaffolds and Dovetail scaffolds revealed a higher concordance compared to the V1 pseudomolecules (**Fig. 5a, b**). At the level of 10-kb fragment alignments, the collinearity between TRITEX positions and mapped position in the Dovetail assembly was 0.982 (Pearson correlation). On average, 94.8 % of aligned sequence in TRITEX/Dovetail scaffold pairs was concordant orientation. We note that the Dovetail assembly was based on the same sequence scaffolds generated from single assembled BAC clones as were used in the Morex V1 pseudomolecule. Hence, the issues of sequence gaps due to failed BAC assemblies and artificial duplication persist. Nevertheless, Dovetail scaffolding did improve the presentation of complete full-length cDNAs by about two percentage points (**Table 4**), most likely by mending occasional sequence breaks within genes.

Third, we compared the TRITEX super-scaffolds to the optical map of the Morex genome constructed by Bionano genome mapping [3, 16]. The optical contigs were aligned to the *in silico* digested TRITEX assembly using Bionano’s Refaligner. Of Nt.BspQ1 sites in the assembly, 95.9 % were covered by high-confidence alignments (score >= 20) of optical contigs and 88.6 % of label sites were aligned. Vice versa, 95.3 % of label sites in the Bionano map were spanned by high-confidence alignments and 90.0 % of Bionano label sites were aligned to the sequence assembly. Label sites covered by alignments, but themselves not aligned (red lines in **Fig. 5c**) may be due to missed label sites in the optical map, gaps in the sequence assembly or alignment uncertainties. We note that it was not possible to align optical contigs to the BAC-based sequence scaffolds as their contiguity is too low (N50: 79 kb, **Table 4**, [34]). In summary, all three comparisons support the high local accuracy of the TRITEX assembly.

To assess the accuracy at the pseudomolecule level, we plotted alignments between chromosomal pseudomolecules of Mascher et al. ([3], Morex V1) and those constructed using TRITEX (Morex V2) and inspected Hi-C contact matrices (**Fig. 6**). The V1 and V2 pseudomolecules were highly collinear. The contact matrices showed the expected Rabl pattern. Several smaller mis-assemblies present in the V1 pseudomolecule were corrected in V2. For example, Morex V1 had a misplaced sequence in the peri-centromeric regions of chromosome 4H (300 – 400 Mb, **Fig. 6c**), which was correctly placed in Morex V2 as supported by the Hi-C contact matrix (**Fig. 6b**).

**Figure 6:**
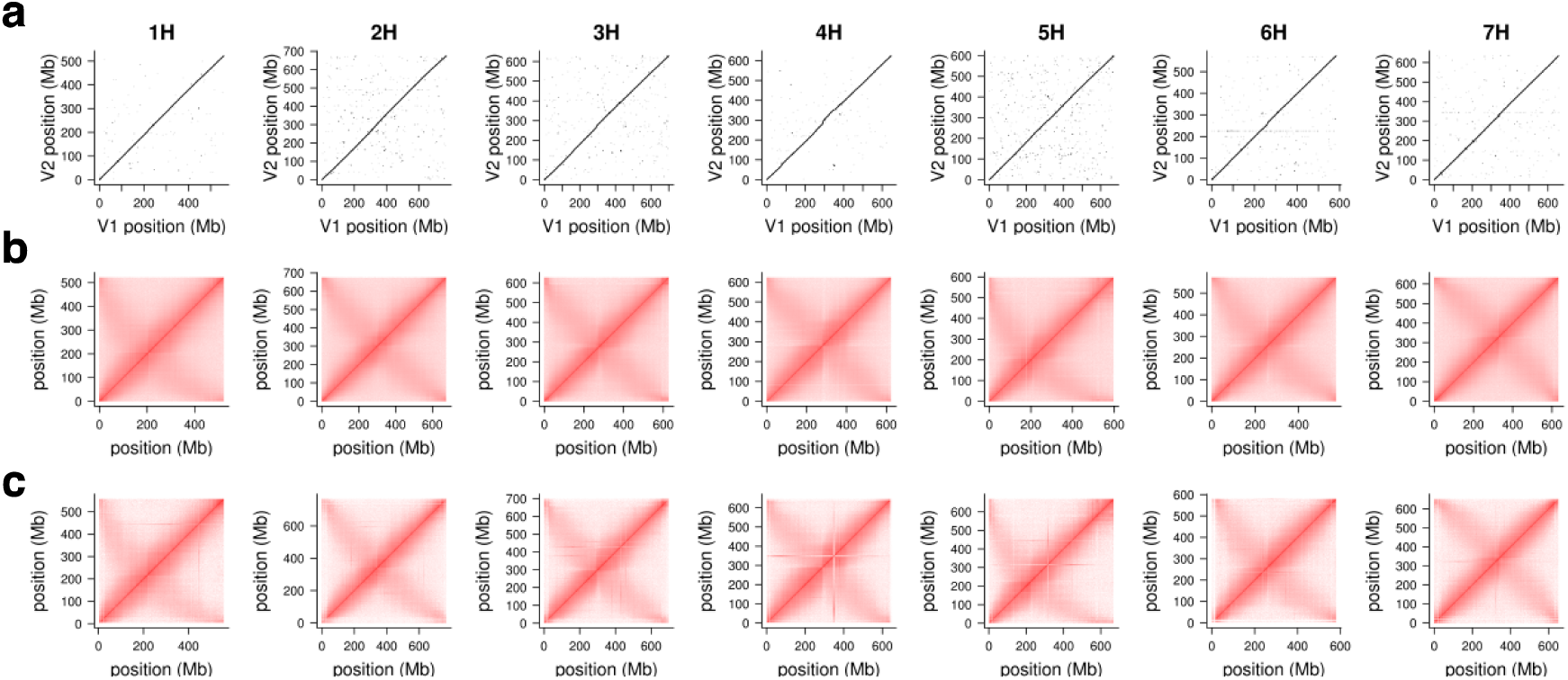
Collinearity of Morex V1 and V2 assemblies. **(a)** Dot plots showing the alignments between the chromosomal pseudomolecules of the Morex V1 and V2 assemblies. **(b)** Intra-chromosomal Hi-C contact matrices of the Morex V2 assembly. **(c)** Intra-chromosomal Hi-C contact matrices of the Morex V1 assembly.

The amounts and characteristics of repetitive sequences such as TEs and tandem repeats represented in sequence assemblies can serve as proxies for assembly quality. We compared the Morex V1 and V2 assemblies according to five criteria: (i) overall TE composition; (ii) presence of highly abundant 20-mers; (iii) amount and localization of tandem repeats; (iv) the amount and age distribution of retrotransposons; and (v) sequence gaps in selected TE families. Overall TE composition was almost identical between the assembly versions (**Table S2**). The chromosomal distribution of highly abundant 20-mers was similar in the Morex V1 and V2 assemblies, but Morex V2 contains more repetitive sequence in peri-centromeric regions (**Fig. S3a**). Similarly, the Morex V2 assembly contains 50 % more tandem repeats than V1. Notably, the number of satellite tandem repeats is almost doubled. Tandem repeats are concentrated in short sequence scaffolds not assigned to chromosomes (chrUn) in Morex V1 and at distal ends of several chromosomes (short arm of 4H, long arms of 4H and 6H) and in peri-centromeric regions (**Fig. S3b**).

The representation of long-terminal repeat (LTR) retrotransposon families was similar in both assemblies (**Table S2**). However, the Morex V2 assembly contains 1,590 (5 %) more intact full-length elements than V1 (**Table S3, Fig. S2a, b**). In both assemblies, the number of retrieved full-length LTRs matches the expectation based on genome size (**Fig. S2c**). Insertion age distributions show that the Morex V2 assembly resolved a higher number of younger Copia elements (**Fig. S2d**). The distinct peak at age 0 in the V1 assembly is most likely caused by a scaffolding artefact from the chromosomal pseudomolecule construction when sequences from the same BAC were arranged in arbitrary order in the Morex V1 pseudomolecules as described above. To understand the impact of sequence gaps on TE representation in the Morex V2 assembly, we performed a similar analysis as for Chinese Spring, using the recently active BARE1 family. The Morex V2 assembly contains more full-length elements than V1 (5,471 vs. 3,469; **Table S1**). Moreover, the percentage of full-length copies that are flanked by a target site duplication (TSD) is higher in the V2 assembly (90% vs. 81%), suggesting fewer chimeric sequences. However, the size distribution of the elements in the Morex V2 assembly indicates a large population of overly large full-length elements (**Table S4**). In contrast, the size distribution of full-length elements in Morex V1 is narrower and shows two characteristic peaks corresponding to the autonomous and non-autonomous subfamilies (T. Wicker, unpublished results). Manual inspection of 50 randomly selected elements between 9,900 and 10,000 bp in length showed that the large sizes of these elements are mainly due to large sequence gaps (i.e. long stretches of N’s). In the 50 manually inspected copies, we found 70 sequence gaps in the internal domain and only 5 short gaps in the LTRs. The latter observation is not surprising as our method to identify full-length copies relied on largely gap-free LTRs. In only three cases, the large size of the element was caused by the genuine insertion of additional TEs. Overall, the Morex V2 assembly had more and larger gaps as TE length increased (**Table S4**), a pattern that is absent from the Morex V1 assembly. In summary, the representation of repetitive sequence is similar in both assembly version of the Morex genome. The longer read lengths and k-mer sizes used in the TRITEX pipeline may have resulted in a better representation of short tandem repeats in V2. However, the gap-free assembly of very recently inserted full-length TEs may benefit from prior complexity reduction such as BAC sequencing.

To facilitate the adoption of the Morex V2 assembly as a common reference sequence by the cereal research community, we annotated the pseudomolecules using the same transcript datasets as used by Mascher et al. [3] for Morex V1, but with an improved version of the PGSB annotation pipeline. A total of 32,787 high-confidence (HC) and 30,871 low-confidence (LC) gene models were annotated on the V2 pseudomolecules. Of the 1,440 BUSCOs (Benchmarking Universal Single-Copy Orthologs, [42]), 98.9 % were completely represented by annotated genes, a 6.4 % increase compared to the V1 annotation. At the same time, the V2 annotation has fewer high-confidence gene models (32,787 [V2] vs. 39,734 [V1]), likely owing to higher assembly contiguity (i.e. fewer fragmented gene models), more stringent thresholds during the annotation process and the incorporation of TE annotations as hints for ab *initio* prediction to reduce the number of transposon-related genes. In a comparison against an independent reference database comprising a curated protein set from 11 grass species, the Morex V1 protein sequences were on average shorter than their V2 counterparts as indicated by a lower alignment coverage. An analysis of sequence gaps in the intergenic space surrounding genes revealed that 90 % of V2, but only 60 % of V1, genes models do not have any “N” bases in their 1-kb flanking regions in the respective sequence assemblies (**Figure S4**). Thus, the Morex V2 gene annotation represents more complete gene models and more regulatory regions around genes compared to the V1 annotation. In conclusion, the TRITEX assembly of Morex constitutes a greatly improved barley reference genome and will serve as an important community resource.

## Discussion

We have developed an open-source pipeline for chromosome-scale sequence assemblies of wheat and barley, and validated its performance by comparison to complementary sequence and mapping resources available for the two species. We believe the main application of TRITEX will be in (i) cereal pan-genomics (i.e. assembling genome sequences for representative genotypes, (ii) phylogenomics (i.e. assembling crop-wild relatives in the Triticeae), and (iii) gene isolation (assisting map-based cloning projects) in the immediate future.

First, in a pan-genomics scenario it is desirable to achieve chromosome-scale sequences of a (few) dozen genotypes representative of major germplasm groups [23]. An open-source pipeline provides the cereal genomics community with a cost-effective platform to generate comparable genome sequences that are amendable to further improvement and refinement. The up-front cost of purchasing hardware (or leasing cloud computing) and (self-) educating researchers in assembly methodology is justified if many assemblies are done. As service fees for assembly may be as high as the expenses for data generation, academic researchers can double the number accessions included in a pan-genomics project if they perform sequence assembly on their own. Alternatives on-site computing infrastructures are national computing infrastructures such as CyVerse [43], de.NBI [44] or SNIC [45].

Second, we anticipate that TRITEX will work well in any diploid or allopolyploid inbreeding Triticeae species. In the Triticeae, important donors of biotic stress resistance loci such as *Ae. sharonensis* [46] or *Ae. longissima* [47] should be amenable to TRITEX assembly. Likewise, our pipeline should be applicable to rye (*Secale cereale*), a minor cereal crop with great importance in East and Central Europe. Although rye is a highly heterozygous, outcrossing species, inbred lines are frequently used by breeders and genomic researchers [48, 49]. TRITEX is likely to work well in species with large and allopolyploid genomes outside the Triticeae tribe such as maize and oats. For maize at least, high-quality sequence assemblies have been constructed with a commercial assembly algorithm [50-52]. However, we must caution users that TRITEX yields assemblies of much lower contiguity and genome representation if closely related haplotypes resident in the same nucleus have to be resolved (Bruno Studer, Martin Mascher; unpublished results). Therefore, we encourage researchers aiming at assembling the genomes of heterozygous, autoploid or dikaryotic species to use long-read sequencing.

Third, Thind et al. [53] recently used chromosomal genomics to assist a gene isolation project. They constructed a megabase-scale sequence scaffold of wheat chromosome 2D of cultivar CH Campala harboring flanking markers of a leaf rust resistance locus to isolate a candidate gene, which was absent from the Chinese Spring reference. In cases, where flow-sorting is not possible, the same purpose (albeit at a higher cost) may be served by TRITEX whole-genome assemblies of the parents of a mapping population or near-isogenic lines carrying mutant introgressions. De novo sequence assembly may be of particular relevance to understanding the molecular basis of plant performance in crop-wild introgression lines derived from wide crosses harboring introgressed segments highly divergent from the reference sequence of the domesticate. For example, these might be pre-breeding material with improved disease resistance, but suffering from linkage drag [54], or released wheat cultivars with alien introgressions conferring superior agronomic performance [55].

The modular layout of our pipeline lends itself to improvement, replacement or simplification of its components. Compared to DeNovoMagic, TRITEX achieves a lower contiguity at the scaffold level and has a higher proportion of internal gaps (**Tables 3, 4**). Frequent gaps and breaks in repetitive sequence are inherent to short-read assemblies. In fact, long-read assemblies with *contig* N50s exceeding *scaffold* N50s of short-read assemblies have been obtained in other plant and animal species [56-58]. Thus, we propose to improve our Illumina-based scaffolding and gap-filling methodology by integrating long-read sequencing into TRITEX in the future. This can be accomplished either by replacing contigging and mate-pair scaffolding entirely with long-read assembly – contingent on the feasibility of obtaining megabase-scale contig N50s in the Triticeae. Alternatively, contig assembly with PE450 reads may be maintained, but long-read sequences could be used for scaffolding and closing gaps. Depending on their accuracy and relative cost-and time-effectiveness, both approaches may be valid for different applications. Long-read sequencing in the Triticeae may adopt the recently developed circular consensus method of Wenger et al. [59] or improved Nanopore sequencing to obtain highly accurate long-reads. These will likely be crucial to resolve homeologs in polyploid wheat where genic sequence divergence between subgenomes is lower than the error-rate of uncorrected long reads.

Our methods for pseudomolecule construction evolved from scripts used for Hi-C mapping of BAC-based sequence scaffolds of barley [3] and whole-genome shotgun assemblies of wheat [4, 17]. They can correct, order and orient along the chromosomes sequence scaffolds of sufficient contiguity and genome representation produced by any sequencing and assembly strategy. We anticipate that even the best long-read assemblies will not be error-free and yield chromosomal contigs without the use complementary linkage information Thus, our methods for assembly correction and super-scaffolding based on linked-read and Hi-C will likely survive the transition to long-reads for contig assembly.

For some research purposes, chromosomal sequences may not be required. If a narrow target interval has been defined in a map-based cloning project, PE450 and MP9 reads may suffice to obtain a single sequence scaffold harboring flanking markers at both sides. In this case, Hi-C and even 10X sequencing can be forgone for faster and cheaper assembly, but at the expense of sequence contiguity.

## Methods

### High molecular weight DNA extraction

High molecular weight (HMW) DNA depleted for plastidal genomes were prepared from fresh leaves of one-week old seedlings of ‘Morex’ using a large-scale phenol:chloroform extraction [17]. In short, the protocol involves isolation of nuclei from fresh leaf material, then the nuclei are treated with proteinase-K, and phenol-chloroform extraction removes protein contamination. Then, the HMW DNA was spooled out of the solution using sodium acetate and ethanol precipitation. The extracted DNA was used for preparation of PCR-free paired-end libraries, mate-pair libraries with specific insert sizes and 10X Chromium libraries.

### Library preparation and sequencing

PCR-free paired-end libraries with 400 – 500 bp insert sizes (PE450) for barley cv. Morex were prepared using a custom protocol using the Illumina Truseq PCR-free library preparation kit. The protocol starts with fragmentation of HMW DNA by ultrasound (Covaris S220, Duty Factor 8%, Peak Incident Power 160, Cycles Per Burst 200, Time 60 seconds) followed by BluePippin size selection on a 1.5% cassette with tight 470bp setting. Then, the size-selected DNA was used as input material for Truseq DNA PCR-free library preparation without the SPB bead-based size selection. The prepared libraries were quantified using the KAPA library quantification kit. Sequencing of the PE450 libraries was done on the HiSeq2500 system in Rapid Run mode (2×266 bp reads). The Morex PE800 paired-end library (insert size range: 700 – 800 kb) as well as MP3 (insert size range: 2-4 kb), MP6 (5-7 kb) and MP9 (8-10 kb) mate-pair libraries were constructed and sequenced (2×150 bp reads) at the University of Illinois Roy. J Carver Biotechnology Center.

To prepare 10X genomics libraries, genomic DNA (gDNA) was quantified by fluorometry (Qubit 2.0). Small fragments (< 40 kb) were removed from ~2 μg of gDNA using pulsed-field electrophoresis on a Blue Pippin instrument (Sage Science, http://www.sagescience.com) following the high-pass protocol. Recovered HMW DNA was evaluated for integrity and size (> 48.5 kb) on a Tapestation 2200 (Agilent, https://www.agilent.com), and quantified (Qubit 2.0, https://www.thermofisher.com/de/en/home/industrial/spectroscopy-elemental-isotope-analysis/molecular-spectroscopy/fluorometers/qubit.html). Library preparation followed the 10X Genome Chromium library protocol v1 (10X Genomics, https://www.10xgenomics.com). Four individual libraries were prepared and uniquely indexed for multiplexing, and quantified by qPCR (Kapa Biosystems). Libraries were normalized and pooled for sequencing on two lanes of an Illumina HiSeq2500 instrument in PE125 mode using v4 chemistry for high output. Pooled libraries were de-multiplexed with Supernova (10X Genomics) and FASTQ files generated with Longranger (10X Genomics).

### Preprocessing of paired-end and mate-pair reads

Overlapping single reads of the PE450 libraries were merged with PEAR [25] (Zavitan), or with BBMerge [26] (Chinese Spring, Morex) using the ‘maxloose’ strictness setting. Error-correction of merged PE450 reads was done with BFC [27] in two passes. After the first BFC pass (correction), reads containing singleton k-mers were trimmed using a k-mer size of 61. Illumina adapters were trimmed from the PE800 read using Cutadapt [60]. Nextera junction adapters and short-insert contaminants were removed from mate-pair reads using NxTrim [61]. Trimmed PE800 and mate-pair reads were corrected with BFC [27] using the hash table of k-mer counts generated from the PE450 reads.

### Unitig assembly

Minia3 ([30], https://github.com/GATB/minia) was used to assemble corrected and trimmed PE450 reads into unitigs. The Minia3 source was assembled to enable k-mer size up to 512 as described in the Minia3 manual. The parameters “-no-bulge-removal -no-tip-removal -no-ec-removal” were used to disable the resolution of ambiguous paths. Iterative Minia3 runs with increasing k-mer sizes (100, 200, 300, 350, 400, 450, 500) were used as proposed in the GATB Minia pipeline (https://github.com/GATB/gatb-minia-pipeline). In the first iteration, the input reads were assembled using a k-mer size of 100. In the subsequent runs, the input reads as well as the assembly of the previous iteration were used as input for the assembler.

### Scaffolding and gap-closing

Error-corrected PE800, MP3, MP6 and MP9 reads were used for scaffolding with SOAPDenovo2 [62]. The “fusion” module (https://github.com/aquaskyline/SOAPdenovo2) was used to prepare the Minia3 unitigs for use with SOAPDenovo2. The “map” was used to align reads to the unitigs. A range of parameters for “pair_num_cutoff” (minimum of read pairs linking two sequences) for each library were tested with the “scaff” module with gap-filling disabled, and the “pair_num_cutoff” value resulting in the best N50 was chosen. Once the best thresholds had been determined, the “scaff” module was run with gap-filling enabled (parameter -F).

GapCloser (https://sourceforge.net/projects/soapdenovo2/files/GapCloser/src/r6/) was used to fill internal gaps in scaffolds using the error-corrected PE450 reads.

### Alignment of 10X Chromium reads and molecule calling

Before alignment, the FASTQ files of read 1 and read 2 were interleaved with Seqtk (https://github.com/lh3/seqtk). Then, the first 23 nt of read 1 were removed and the instrument name in the Illumina read identifier was replaced by the 10X barcode (the first 16 nt of read 1). Illumina adapters were trimmed using cutadapt [60] using the adapter sequence AGATCGGAAGAGC. Reads shorter than 30 bp after trimming were discarded. Trimmed reads were aligned to the assembly after the GapCloser step using Minimap2 [36]. Alignment records were converted to the Binary Sequence Alignment/Map (BAM) Format with SAMtools [63]. BAM files were sorted twice with Novosort (http://www.novocraft.com/products/novosort/). First, reads were sorted by alignment positions and duplicated read pairs were flagged. Then, reads were sorted by name to group read pairs with identical barcodes together. After sorting, BAM files were converted into BEDPE format using BEDTools [64]. Duplicated and supplementary alignments were ignored. Only read pairs with a mapping quality >= 20 were retained. Read pairs with an estimated insert size above 800 bp were discarded. Read pairs were written into a table with four columns (chromosome, start, end, barcode), which was sorted by barcode and alignment position using GNU sort. Read pairs having the same barcode and mapping within 500 kb of each other were assigned to the same molecule. A table recording the barcodes as well as the start and end points of each molecule was exported as a text file. Molecules shorter than 1 kb were discarded.

### 10X super-scaffolding

A graph structure is constructed in which nodes are scaffolds and edges are drawn between if a sufficient number of 10X links meet certain criteria. Only molecules mapping within 100 kb of the scaffold ends are considered. Edges between scaffolds are accepted only if they are supported by molecules from more than one Chromium library. In the initial graph, only molecules connecting scaffolds anchored by POPSEQ to the same chromosome within 5 cM of each other are allowed. A minimum spanning tree is computed with functionalities of the R package igraph [65]. Subsequently, heuristics are applied to resolve branches to obtain subgraphs that are paths. Heuristics include (i) the removal of tips (nodes of rank one), (ii) nodes corresponding to small (< 10 kb) scaffolds, and (iii) cutting the tree at branch points by removing edges. These subgraphs are the initial super-scaffolds and the order of scaffolds in them is determined by the (well-defined) path traversing each subgraph. The orientation of a scaffold S within a super-scaffold X is determined by calculating the mean position of molecules linking S to other scaffolds in X up to five bins upstream (to the left) or downstream (to the right) of S. If the average position of downstream links is larger than that of the upstream links, the orientation of S is “forward”, and “reverse” otherwise. Super-scaffolds are assigned to POPSEQ genetic positions by lifting positional information from their constituent scaffolds (see below) and computing a consensus. Super-scaffold construction is repeated with different thresholds for the minimum number of read pairs to assign molecules to scaffold (2 to 10) and the minimum number of molecules required to an accept edges between scaffold pairs (2 to 10) and the assembly with the highest N50 is selected for further steps.

### Hi-C map construction

Scaffolds after the GapCloser step were digested *in silico* with EMBOSS restrict [66]. Reads were aligned to the scaffolds and assigned to restriction fragments as described by Beier et al. [34]. In constrast to Beier et al., we used Minimap2 [36] instead of BWA-MEM [67] for read alignment. Scaffolds were assigned to chromosomes using the POPSEQ genetic maps of wheat and barley [12, 13] as described by Avni et al. [17]. POPSEQ marker sequences (the scaffolds of synthetic wheat W7984 assembled by Chapman et al. [13] or the WGS contigs of the International Barley Sequencing Consortium (2012) [39]) were aligned to the scaffolds using Minimap2 [36]. POPSEQ positional information and Hi-C links were lifted from scaffolds to super-scaffolds. Super-scaffolds were ordered and oriented as described by Beier et al. [34]. Intra-chromosomal Hi-C matrices were normalized with HiCNorm [68] and visually inspected in a locally installed R Shiny app. The code for the Shiny app is provided in the Bitbucket repository.

### Discovery and correction of mis-assemblies

We follow the approach of Putnam et al. [69] and Ghurye et al. [33] by detecting sequence mis-joins made during the scaffold stage by inspecting the physical coverage with either 10X molecules or Hi-C links. To find breakpoints based on 10X (Hi-C) data, scaffolds were divided into 200 bp (1 kb) bins and the number of molecules (Hi-C links) spanning each bin was calculated. Links counts were normalized by distance from the scaffolds ends. Drops in 10X coverage below one eighth of the genome-wide average were considered as breakpoints. The minimum distance between two breakpoints was 50 kb. After breaking chimeras, POPSEQ marker positions, 10X molecule boundaries and Hi-C links were lifted to the corrected assemblies. This procedure was repeated until no breakpoints were detected. Drops in Hi-C coverage below one sixteenth of the genome-wide average were considered as potential breakpoints and diagnostic plots summarizing POPSEQ marker information as well as Hi-C coverage along scaffolds were visually inspected. If necessary, breakpoint coordinates were adjusted manually.

### Dovetail assembly

Chicago libraries were prepared from leaves of barley cv. Morex by Dovetail Inc. as described by Putnam et al. [41]. The HiRise algorithm was used to scaffold the non-redundant BAC-based sequence contigs assembled by Mascher et al. [3] (accessible from http://dx.doi.org/10.5447/IPK/2016/30).

### Optical map alignment

The genome-wide optical map of barley cv. Morex [3] was retrieved from http://dx.doi.org/10.5447/IPK/2016/31 and aligned to the TRITEX super-scaffolds using Bionano RefAligner (https://bionanogenomics.com). Custom scripts were used for importing alignments into R and visualization (https://bitbucket.org/tritexassembly/tritexassembly.bitbucket.io/src/master/miscellaneous/bionano.R).

### Assembly-to-assembly alignment

Recent versions of Minimap2 ([36], https://github.com/lh3/minimap2) were used for assembly-to-assembly alignment. Alignment records were written to PAF format and imported into R for visualization and calculation of summary statistics.

### Transcript alignment

Transcript datasets were aligned with GMAP [70] version 2018-07-04 to genomic references. Alignment records were written to GFF files, from which coverage and alignment identity for mRNA alignments were extracted.

### Barley gene annotation

Our annotation pipeline combines three types of evidence for structural gene annotation in plants: protein homology, expression data and *ab initio* prediction. For homology-based annotation, we combined available Triticeae protein sequences obtained from UniProt (05/10/2016). These protein sequences were mapped to the nucleotide sequence of the Morex V2 pseudomolecules using the splice-aware alignment software GenomeThreader [71] (version 1.7.1; arguments -startcodon -finalstopcodon -species rice -gcmincoverage 70 - prseedlength 7 -prhdist 4). Full-length cDNA [72] and IsoSeq [3] nucleotide sequences were aligned to the Morex V2 pseudomolecules using GMAP [70] (version 2018-07-04, standard parameters). Illumina RNA-seq datasets were first mapped using Hisat2 [73] (version 2.0.4, parameter --dta) and subsequently assembled into transcript sequences by Stringtie [74] (version 1.2.3, parameters -m 150 -t -f 0.3. Full-length cDNA, IsoSeq sequences and RNASeq datasets are described in [3]. All transcripts from flcDNA, IsoSeq and RNASeq were combined using Cuffcompare [75] (version 2.2.1) and subsequently merged with Stringtie (version 1.2.3, parameters --merge -m 150) to remove fragments and redundant structures. Next, we used Transdecoder (version 3.0.0, https://github.com/TransDecoder) to find potential open reading frames and to predict protein sequences. We used BLASTP [76] (ncbi-blast-2.3.0+, parameters -max_target_seqs 1 -evalue 1e-05) to compare potential protein sequences with a trusted set of reference proteins (Uniprot Magnoliophyta, reviewed/Swissprot, downloaded on 3 Aug 2016) and used hmmscan [77] (version 3.1b2) to identify conserved protein family domains for all potential proteins. BLAST and hmmscan results were fed back into Transdecoder-predict to select the best translations per transcript sequence.

An independent *ab initio* annotation using Augustus [78] (version 3.3.2) was carried out to further improve structural gene annotation. To minimize over-prediction, hint files using the above mentioned IsoSeq, flcDNA, RNASeq, protein evidences and TE predictions were generated. The wheat model was used for prediction.

All structural gene annotations were joined by feeding them into EVidenceModeler [79] and weights were adjusted according to the input source. To refine gene models, we also incorporated the Barley reference transcript database (BaRT) as an additional source. All BaRT transcripts were aligned to the new Morex V2 assembly using GMAP (version 2018-07-04) and output was converted into GFF format and subsequently fed into EVidenceModeler. Finally, redundant protein sequences were removed to form a single non-redundant candidate dataset. To categorize candidates into complete and valid genes, non-coding transcripts, pseudogenes and transposable elements, we applied a confidence classification protocol. Candidate protein sequences were compared against the following three manually curated databases using BLAST: first, PTREP, a database of hypothetical proteins that contains deduced amino acid sequences in which, in many cases, frameshifts have been removed, which is useful for the identification of divergent TEs having no significant similarity at the DNA level; second, UniPoa, a database comprised of annotated Poaceae proteins; third, UniMag, a database of validated magnoliophyta proteins. UniPoa and UniMag protein sequences were downloaded from Uniprot and further filtered for complete sequences with start and stop codons. Best hits were selected for each predicted protein to each of the three databases. Only hits with an E-value below 10^-10^ were considered.

Furthermore, only hits with subject coverage (for protein references) or query coverage (transposon database) above 75% were considered significant and protein sequences were further classified using the following confidence: a high confidence (HC) protein sequence is complete and has a subject and query coverage above the threshold in the UniMag database (HC1) or no blast hit in UniMag but in UniPoa and not TREP (HC2); a low confidence (LC) protein sequence is not complete and has a hit in the UniMag or UniPoa database but not in TREP (LC1), or no hit in UniMag and UniPoa and TREP but the protein sequence is complete. The tag REP was assigned for protein sequences not in UniMag and complete but with hits in TREP.

Functional annotation of predicted protein sequences was done using the AHRD pipeline (https://github.com/groupschoof/AHRD). Completeness of the predicted gene space was measured with BUSCO [42] (version 3.02, orthodb9).

### Analysis of 5’ and 3’ flanking regions of gene models

5’ and 3’ flanking nucleotide sequences of increasing lengths in the range from 1 kb to 10 kb upstream and downstream of predicted gene models were extracted and sequences containing N were discarded. The remaining N-free sequences were counted and plotted as a percentage of the total number of predicted gene models.

### Repeat annotation

Transposons where detected and classified by an homology search against the REdat_9.7_Triticeae subset of the PGSB transposon library [80] using vmatch (http://www.vmatch.de) using following parameter settings: identity >= 70%, minimal hit length 75 bp, seed length 12 bp (exact command line: -d -p -l 75 -identity 70 -seedlength 12 - exdrop 5). The vmatch output was filtered for redundant hits via a priority-based approach, which assigns higher scoring matches first and either shortens (< 90% coverage and >= 50bp rest length) or removes lower scoring overlaps, leading to an overlap free annotation.

Full-length LTR retrotransposons where identified with LTRharvest [81] using the following parameters: “overlaps best -seed 30 -minlenltr 100 -maxlenltr 2000 -mindistltr 3000 - maxdistltr 25000 -similar 85 -mintsd 4 -maxtsd 20 -motif tgca -motifmis 1 -vic 60 -xdrop 5 - mat 2 -mis -2 -ins -3 -del -3”. All candidates from the LTRharvest output were subsequently annotated for PfamA domains using hmmer3 [82] and stringently filtered for false positives by several criteria, the main ones being a lack of at least one typical retrotransposon domain [e.g. reverse transcriptase (RT), RNase H (RH), integrase (INT), protease (PR), etc.)] and a tandem repeat content > 25%. The inner domain order served as a criterion for the classification into the Gypsy (RT-RH-INT) or Copia (INT-RT-RH) superfamily abbreviated as RLG and RLC. Elements missing either INT or RT were classified as RLX. The insertion age of each full-length LTR retrotransposon was estimated based on the accumulated divergence between its 5’ and 3’ long terminal repeats and a mutation rate of 1.3 × 10^-8^ [83].

Tandem repeats where identified with the TandemRepeatFinder [84] using default parameters and subjected to an overlap removal as decribed above, prioritizing longer and higher scoring elements. K-mer frequencies were calculated with Tallymer [85].

### Representation of selected TE families

We identified full-length copies belonging to single TE families in the assemblies of Chinese Spring and Morex. Our pipeline uses BLASTN [76] to search for long terminal repeats (LTRs) that occur at a user-defined distance range in the same orientation. For RLC_BARE1 and RLC_Angela elements, the two LTRs had to be found within a range of 7,800-9,300 bp (a consensus RLC_BARE1 sequence has a length of approximately 8,700 bp), while a range from 6,000-10,000 bp was allowed for RLG_Sabrina elements. Multiple different LTR consensus sequences were used for the searches in order to cover the intra-family diversity. Five LTR consensus sequences each were used for RLC_BARE1 and RLG_Sabrina elements, while 18 LTR consensus sequences were used for RLC_Angela elements (to cover the much wider diversity of this family in the three wheat subgenomes). The LTR consensus sequences from the same families are 73-92% identical to each other, reflecting the considerable intra-family diversity.

For the current analysis, full-length copies of RLC_BARE1 and their wheat homologs RLC_Angela elements were extracted because these are the most abundant families in barley and wheat genomes, respectively. RLG_Sabrina was chosen because preliminary analyses showed that this TE family has not been active in wheat for a long time and thus is represented mostly by old copies.

To validate the extracted TE populations, the size range of all isolated copies as well as the number of copies that are flanked by a target site duplication (TSD) were determined. A TSD is accepted if it contains at least 3 matches between 5’ and 3’ TSD (e.g. ATGCG and ACGAG). This low stringency was applied because our previous study showed that TSD generation is error-prone, and thus multiple mismatches can be expected [86]. In a survey, 80-90% of all isolated full-length elements were flanked by a TSD.

Our pipeline also extracts so-called “solo-LTRs”. These are products of intra-element recombination that results in the loss of the internal domain and the generation of a chimeric solo-LTR sequence. Solo-LTRs were used as a metric of how well short repetitive sequences are assembled.

### Data availability

Paired-end and mate-pair data for Zavitan [17] and Chinese Spring [4] were retrieved from EMBL ENA (accession numbers: PRJEB31422 and SRP114784). Paired-end, mate-pair, 10× and Chicago data generated for Morex were deposited under ENA project ID PRJEB31444. Hi-C data for barley cv. Morex [3] are available under ENA accession PRJEB14169. Sequence assemblies generated in the present study are accessible under the following Digital Object Identifiers (DOIs) in the Plant Genomics & Phenomics Research Data Repository [87]: doi:10.5447/IPK/2019/6 (wheat assemblies); doi:10.5447/IPK/2019/8 (barley Morex V2 assembly); doi: 10.5447/IPK/2019/7 (barley Dovetail assembly). DOIs were registered with e!DAL [88].

### Access to source code, documentation and software versions

Z shell scripts were used to call assembly and alignment software. GNU Parallel [89] was used for parallel computing. Scaffolding with 10X and Hi-C data was implemented in GNU AWK (https://www.gnu.org/s/gawk/manual/gawk.html) and R [31] scripts. R code relies heavily on the data.table package (https://cran.r-project.org/package=data.table) for in-memory data management and analysis. A step-by-step usage guide was prepared using the AsciiDoc markup language (https://asciidoctor.org/docs/what-is-asciidoc/) is available from https://nomagicassembly.bitbucket.io. A list of software versions known to work with TRITEX is included in the usage guide. Source code is hosted in a public Bitbucket repository (https://bitbucket.org/nomagicassembly/nomagicassembly.bitbucket.io). Assemblies were run on compute servers at IPK Gatersleben. The most powerful of these has 72 physical cores (Intel Xeon E7-8890 v3) and 2 TB of main memory. We had access to 100 TB of hard disk space and 22 TB of SSD storage.

## Author Contributions

M.M. and N.S. conceived the study. S.P. and A.H. performed library preparation and sequencing. M.M. designed the computational workflow. C.M. and M.M. performed assembly. T.W. and H.G. analyzed transposable elements. T.L., H.G., M.S. and K.F.X.M. performed genome annotation. J.E. and C.P. constructed 10X libraries. U.S. supervised IT administration. I.B., C.L., M.M., K.F.X.M, G.J.M, A.H.S, N.S. and R.W. contributed the Dovetail assembly. C.M. and M.M. wrote the paper with input from all co-authors.

## Acknowledgements

This research was supported by grants from the German Federal Ministry of Education and Research to N.S., M.M., U.S., M.S. and K.F.X.M (‘SHAPE’, FKZ 031B0190), to U.S. and K.F.X.M (‘De.NBI’, FKZ 031A536) and from the German Federal Ministry of Food and Agriculture to K.F.X.M (grant 2819103915, ‘Wheatseq’). Support for 10X sequencing was provided by a research grant from Genome Canada and Genome Prairie (to C.P. and J.E). Tim Close (University of California, Riverside) contributed funds from Hatch Project CA-R-BPS-5306-H to the Dovetail assembly. We are indebted to Rayan Chikhi for pointing us to the multi k-mer approach. We are grateful to Manuela Knauft, Ines Walde and Susanne König for technical assistance. We thank Anne Fiebig for handling data submission and Jens Bauernfeind, Thomas Münch and Heiko Miehe for IT administration. We thank Johannes Heilmann for introducing us to AsciiDoc, and Sara Giulia Milner, Kevin Koh, Liangliang Gao, Juan Gutierrez Gonzalez, Burkhard Steuernagel, Kumar Gaurav and Raz Avni for feedback on previous versions of the workflow.

## References

1. Schulte D, Close TJ, Graner A, Langridge P, Matsumoto T, Muehlbauer G, Sato K, Schulman AH, Waugh R, Wise RP: The international barley sequencing consortium— at the threshold of efficient access to the barley genome. Plant physiology 2009, 149:142–147.

2. Gill BS, Appels R, Botha-Oberholster AM, Buell CR, Bennetzen JL, Chalhoub B, Chumley F, Dvorak J, Iwanaga M, Keller B, et al: A workshop report on wheat genome sequencing: International Genome Research on Wheat Consortium. Genetics 2004, 168:1087–1096.

3. Mascher M, Gundlach H, Himmelbach A, Beier S, Twardziok SO, Wicker T, Radchuk V, Dockter C, Hedley PE, Russell J, et al: A chromosome conformation capture ordered sequence of the barley genome. Nature 2017, 544:427–433.

4. The International Wheat Genome Sequencing Consortium (IWGSC): Shifting the limits in wheat research and breeding using a fully annotated reference genome. Science 2018, 361:eaar7191.

5. Maccaferri M, Harris NS, Twardziok SO, Pasam RK, Gundlach H, Spannagl M, Ormanbekova D, Lux T, Prade VM, Milner SG, et al: Durum wheat genome highlights past domestication signatures and future improvement targets. Nature Genetics 2019.

6. Luo MC, Gu YQ, Puiu D, Wang H, Twardziok SO, Deal KR, Huo N, Zhu T, Wang L, Wang Y, et al: Genome sequence of the progenitor of the wheat D genome Aegilops tauschii. Nature 2017, 551:498–502.

7. Ling HQ, Ma B, Shi X, Liu H, Dong L, Sun H, Cao Y, Gao Q, Zheng S, Li Y, et al: Genome sequence of the progenitor of wheat A subgenome Triticum urartu. Nature 2018, 557:424–428.

8. Avni R, Nave M, Barad O, Baruch K, Twardziok SO, Gundlach H, Hale I, Mascher M, Spannagl M, Wiebe K: Wild emmer genome architecture and diversity elucidate wheat evolution and domestication. Science 2017, 357:93–97.

9. McPherson JD, Marra M, Hillier L, Waterston RH, Chinwalla A, Wallis J, Sekhon M, Wylie K, Mardis ER, Wilson RK, et al: A physical map of the human genome. Nature 2001, 409:934–941.

10. Beier S, Himmelbach A, Schmutzer T, Felder M, Taudien S, Mayer KF, Platzer M, Stein N, Scholz U, Mascher M: Multiplex sequencing of bacterial artificial chromosomes for assembling complex plant genomes. Plant biotechnology journal 2016, 14:1511–1522.

11. Choulet F, Alberti A, Theil S, Glover N, Barbe V, Daron J, Pingault L, Sourdille P, Couloux A, Paux E: Structural and functional partitioning of bread wheat chromosome 3B. Science 2014, 345:1249721.

12. Mascher M, Muehlbauer GJ, Rokhsar DS, Chapman J, Schmutz J, Barry K, Muñoz-Amatriaín M, Close TJ, Wise RP, Schulman AH, et al: Anchoring and ordering NGS contig assemblies by population sequencing (POPSEQ). The Plant Journal 2013, 76:718–727.

13. Chapman JA, Mascher M, Buluc A, Barry K, Georganas E, Session A, Strnadova V, Jenkins J, Sehgal S, Oliker L, et al: A whole-genome shotgun approach for assembling and anchoring the hexaploid bread wheat genome. Genome Biol 2015, 16:26.

14. Burton JN, Adey A, Patwardhan RP, Qiu R, Kitzman JO, Shendure J: Chromosomescale scaffolding of de novo genome assemblies based on chromatin interactions. Nature biotechnology 2013, 31:1119.

15. Kaplan N, Dekker J: High-throughput genome scaffolding from in vivo DNA interaction frequency. Nature biotechnology 2013, 31:1143.

16. Lam ET, Hastie A, Lin C, Ehrlich D, Das SK, Austin MD, Deshpande P, Cao H, Nagarajan N, Xiao M: Genome mapping on nanochannel arrays for structural variation analysis and sequence assembly. Nature biotechnology 2012, 30:771.

17. Avni R, Nave M, Barad O, Baruch K, Twardziok SO, Gundlach H, Hale I, Mascher M, Spannagl M, Wiebe K, et al: Wild emmer genome architecture and diversity elucidate wheat evolution and domestication. Science 2017, 357:93–97.

18. Zhu T, Wang L, Rodriguez JC, Deal KR, Avni R, Distelfeld A, McGuire PE, Dvorak J, Luo MC: Improved Genome Sequence of Wild Emmer Wheat Zavitan with the Aid of Optical Maps. G3 (Bethesda) 2019, 9:619–624.

19. Callaway E: Small group scoops international effort to sequence huge wheat genome. Nature News 2017.

20. Clavijo BJ, Venturini L, Schudoma C, Accinelli GG, Kaithakottil G, Wright J, Borrill P, Kettleborough G, Heavens D, Chapman H: An improved assembly and annotation of the allohexaploid wheat genome identifies complete families of agronomic genes and provides genomic evidence for chromosomal translocations. Genome research 2017, 27:885–896.

21. Zimin AV, Puiu D, Luo MC, Zhu T, Koren S, Marcais G, Yorke JA, Dvorak J, Salzberg SL: Hybrid assembly of the large and highly repetitive genome of Aegilops tauschii, a progenitor of bread wheat, with the MaSuRCA mega-reads algorithm. Genome Res 2017, 27:787–792.

22. Zimin AV, Puiu D, Hall R, Kingan S, Clavijo BJ, Salzberg SL: The first near-complete assembly of the hexaploid bread wheat genome, Triticum aestivum. Gigascience 2017.

23. Monat C, Schreiber M, Stein N, Mascher M: Prospects of pan-genomics in barley. Theoretical and Applied Genetics 2018:1–12.

24. International Wheat Genome Sequencing Consortium: Shifting the limits in wheat research and breeding using a fully annotated reference genome. Science 2018, 361:eaar7191.

25. Zhang J, Kobert K, Flouri T, Stamatakis A: PEAR: a fast and accurate Illumina Paired-End reAd mergeR. Bioinformatics 2014, 30:614–620.

26. Bushnell B, Rood J, Singer E: BBMerge - Accurate paired shotgun read merging via overlap. PLoS One 2017, 12:e0185056.

27. Li H: BFC: correcting Illumina sequencing errors. Bioinformatics 2015, 31:2885–2887.

28. Melsted P, Halldorsson BV: KmerStream: streaming algorithms for k-mer abundance estimation. Bioinformatics 2014, 30:3541–3547.

29. Chikhi R, Limasset A, Medvedev P: Compacting de Bruijn graphs from sequencing data quickly and in low memory. Bioinformatics 2016, 32:i201–i208.

30. Chikhi R, Rizk G: Space-efficient and exact de Bruijn graph representation based on a Bloom filter. Algorithms Mol Biol 2013, 8:22.

31. Team RC: R: A language and environment for statistical computing. R Foundation for Statistical Computing, Vienna, Austria. 2016. 2017.

32. Sahlin K, Chikhi R, Arvestad L: Assembly scaffolding with PE-contaminated mate-pair libraries. Bioinformatics 2016, 32:1925–1932.

33. Ghurye J, Pop M, Koren S, Bickhart D, Chin CS: Scaffolding of long read assemblies using long range contact information. BMC Genomics 2017, 18:527.

34. Beier S, Himmelbach A, Colmsee C, Zhang X-Q, Barrero RA, Zhang Q, Li L, Bayer M, Bolser D, Taudien S, et al: Construction of a map-based reference genome sequence for barley, Hordeum vulgare L. Scientific Data 2017, 4:170044.

35. Lu F-H, McKenzie N, Kettleborough G, Heavens D, Clark MD, Bevan MW: Independent assessment and improvement of wheat genome sequence assemblies using Fosill jumping libraries. GigaScience 2018, 7:giy053.

36. Li H: Minimap2: pairwise alignment for nucleotide sequences. Bioinformatics 2018, 1:7.

37. Mochida K, Yoshida T, Sakurai T, Ogihara Y, Shinozaki K: TriFLDB: a database of clustered full-length coding sequences from Triticeae with applications to comparative grass genomics. Plant Physiol 2009, 150:1135–1146.

38. Wicker T, Sabot F, Hua-Van A, Bennetzen JL, Capy P, Chalhoub B, Flavell A, Leroy P, Morgante M, Panaud O, et al: A unified classification system for eukaryotic transposable elements. Nature Reviews Genetics 2007, 8:973.

39. International Barley Genome Sequencing Consortium: A physical, genetic and functional sequence assembly of the barley genome. Nature 2012, 491:711.

40. Schnable PS, Ware D, Fulton RS, Stein JC, Wei F, Pasternak S, Liang C, Zhang J, Fulton L, Graves TA: The B73 maize genome: complexity, diversity, and dynamics. science 2009, 326:1112–1115.

41. Putnam NH, O’Connell BL, Stites JC, Rice BJ, Blanchette M, Calef R, Troll CJ, Fields A, Hartley PD, Sugnet CW: Chromosome-scale shotgun assembly using an in vitro method for long-range linkage. Genome research 2016.

42. Simao FA, Waterhouse RM, Ioannidis P, Kriventseva EV, Zdobnov EM: BUSCO: assessing genome assembly and annotation completeness with single-copy orthologs. Bioinformatics 2015, 31:3210–3212.

43. Joyce BL, Haug-Baltzell AK, Hulvey JP, McCarthy F, Devisetty UK, Lyons E: Leveraging CyVerse Resources for De Novo Comparative Transcriptomics of Underserved (Non-model) Organisms. J Vis Exp 2017.

44. Schmutzer T, Bolger ME, Rudd S, Chen J, Gundlach H, Arend D, Oppermann M, Weise S, Lange M, Spannagl M, et al: Bioinformatics in the plant genomic and phenomic domain: The German contribution to resources, services and perspectives. J Biotechnol 2017, 261:37–45.

45. Toor S, Lindberg M, Falman I, Vallin A, Mohill O, Freyhult P, Nilsson L, Agback M, Viklund L, Zazzik H, et al: SNIC Science Cloud (SSC): A National-Scale Cloud Infrastructure for Swedish Academia. In 2017 IEEE 13th International Conference on e-Science (e-Science); 24-27 Oct. 2017. 2017: 219–227.

46. Yu G, Champouret N, Steuernagel B, Olivera PD, Simmons J, Williams C, Johnson R, Moscou MJ, Hernandez-Pinzon I, Green P, et al: Discovery and characterization of two new stem rust resistance genes in Aegilops sharonensis. Theor Appl Genet 2017, 130:1207–1222.

47. Huang S, Steffenson BJ, Sela H, Stinebaugh K: Resistance of Aegilops longissima to the Rusts of Wheat. Plant Dis 2018, 102:1124–1135.

48. Geiger H, Miedaner T: Rye breeding. Cereals 2009, 3:157–181.

49. Bauer E, Schmutzer T, Barilar I, Mascher M, Gundlach H, Martis MM, Twardziok SO, Hackauf B, Gordillo A, Wilde P: Towards a whole-genome sequence for rye (Secale cereale L.). The Plant Journal 2017, 89:853–869.

50. Hirsch C, Hirsch CD, Brohammer AB, Bowman MJ, Soifer I, Barad O, Shem-Tov D, Baruch K, Lu F, Hernandez AG: Draft assembly of elite inbred line PH207 provides insights into genomic and transcriptome diversity in maize. The Plant Cell 2016:tpc. 00353.02016.

51. Springer NM, Anderson SN, Andorf CM, Ahern KR, Bai F, Barad O, Barbazuk WB, Bass HW, Baruch K, Ben-Zvi G, et al: The maize W22 genome provides a foundation for functional genomics and transposon biology. Nat Genet 2018, 50:1282–1288.

52. Unterseer S, Seidel MA, Bauer E, Haberer G, Hochholdinger F, Opitz N, Marcon C, Baruch K, Spannagl M, Mayer KFX, Schön C-C: European Flint reference sequences complement the maize pan-genome. bioRxiv 2017:103747.

53. Thind AK, Wicker T, Simkova H, Fossati D, Moullet O, Brabant C, Vrana J, Dolezel J, Krattinger SG: Rapid cloning of genes in hexaploid wheat using cultivar-specific long-range chromosome assembly. Nat Biotechnol 2017, 35:793–796.

54. Wendler N, Mascher M, Himmelbach A, Johnston P, Pickering R, Stein N: Bulbosum to go: a toolbox to utilize Hordeum vulgare/bulbosum introgressions for breeding and beyond. Molecular plant 2015, 8:1507–1519.

55. Xue S, Kolmer JA, Wang S, Yan L: Mapping of Leaf Rust Resistance Genes and Molecular Characterization of the 2NS/2AS Translocation in the Wheat Cultivar Jagger. G3 (Bethesda) 2018, 8:2059–2065.

56. Jiao Y, Peluso P, Shi J, Liang T, Stitzer MC, Wang B, Campbell MS, Stein JC, Wei X, Chin CS, et al: Improved maize reference genome with single-molecule technologies. Nature 2017, 546:524–527.

57. Schmidt MH, Vogel A, Denton AK, Istace B, Wormit A, van de Geest H, Bolger ME, Alseekh S, Mass J, Pfaff C, et al: De Novo Assembly of a New Solanum pennellii Accession Using Nanopore Sequencing. Plant Cell 2017, 29:2336–2348.

58. Wang M, Tu L, Yuan D, Zhu Shen C, Li J, Liu F, Pei L, Wang P, Zhao G, et al: Reference genome sequences of two cultivated allotetraploid cottons, Gossypium hirsutum and Gossypium barbadense. Nat Genet 2019, 51:224–229.

59. Wenger AM, Peluso P, Rowell WJ, Chang P-C, Hall RJ, Concepcion GT, Ebler J, Fungtammasan A, Kolesnikov A, Olson ND, et al: Highly-accurate long-read sequencing improves variant detection and assembly of a human genome. bioRxiv 2019:519025.

60. Martin M: Cutadapt removes adapter sequences from high-throughput sequencing reads. EMBnet Journal 2011, 17:pp. 10–12.

61. O’Connell J, Schulz-Trieglaff O, Carlson E, Hims MM, Gormley NA, Cox AJ: NxTrim: optimized trimming of Illumina mate pair reads. Bioinformatics 2015, 31:2035–2037.

62. Luo R, Liu B, Xie Y, Li Z, Huang W, Yuan J, He G, Chen Y, Pan Q, Liu Y: SOAPdenovo2: an empirically improved memory-efficient short-read de novo assembler. Gigascience 2012, 1:18.

63. Li H, Handsaker B, Wysoker A, Fennell T, Ruan J, Homer N, Marth G, Abecasis G, Durbin R: The sequence alignment/map format and SAMtools. Bioinformatics 2009, 25:2078–2079.

64. Quinlan AR, Hall IM: BEDTools: a flexible suite of utilities for comparing genomic features. Bioinformatics 2010, 26:841–842.

65. Csardi G, Nepusz T: The igraph software package for complex network research. InterJournal, Complex Systems 2006, 1695:1–9.

66. Rice P, Longden I, Bleasby A: EMBOSS: the European molecular biology open software suite. Trends in genetics 2000, 16:276–277.

67. Li H: Aligning sequence reads, clone sequences and assembly contigs with BWA-MEM. arXiv preprint arXiv:13033997 2013.

68. Hu M, Deng K, Selvaraj S, Qin Z, Ren B, Liu JS: HiCNorm: removing biases in Hi-C data via Poisson regression. Bioinformatics 2012, 28:3131–3133.

69. Putnam NH, O’Connell BL, Stites JC, Rice BJ, Blanchette M, Calef R, Troll CJ, Fields A, Hartley PD, Sugnet CW, et al: Chromosome-scale shotgun assembly using an in vitro method for long-range linkage. Genome Res 2016, 26:342–350.

70. Wu TD, Watanabe CK: GMAP: a genomic mapping and alignment program for mRNA and EST sequences. Bioinformatics 2005, 21:1859–1875.

71. Gremme G, Brendel V, Sparks ME, Kurtz S: Engineering a software tool for gene structure prediction in higher organisms. Information and Software Technology 2005, 47:965–978.

72. Matsumoto T, Tanaka T, Sakai H, Amano N, Kanamori H, Kurita K, Kikuta A, Kamiya K, Yamamoto M, Ikawa H, et al: Comprehensive sequence analysis of 24,783 barley full-length cDNAs derived from 12 clone libraries. Plant Physiol 2011, 156:20–28.

73. Kim D, Langmead B, Salzberg SL: HISAT: a fast spliced aligner with low memory requirements. Nat Methods 2015, 12:357–360.

74. Pertea M, Pertea GM, Antonescu CM, Chang TC, Mendell JT, Salzberg SL: StringTie enables improved reconstruction of a transcriptome from RNA-seq reads. Nat Biotechnol 2015, 33:290–295.

75. Ghosh S, Chan CK: Analysis of RNA-Seq Data Using TopHat and Cufflinks. Methods Mol Biol 2016, 1374:339–361.

76. Altschul SF, Gish W, Miller W, Myers EW, Lipman DJ: Basic local alignment search tool. J Mol Biol 1990, 215:403–410.

77. Eddy SR: Accelerated Profile HMM Searches. PLoS Comput Biol 2011, 7:e1002195.

78. Stanke M, Schoffmann O, Morgenstern B, Waack S: Gene prediction in eukaryotes with a generalized hidden Markov model that uses hints from external sources. BMC Bioinformatics 2006, 7:62.

79. Haas BJ, Salzberg SL, Zhu W, Pertea M, Allen JE, Orvis J, White O, Buell CR, Wortman JR: Automated eukaryotic gene structure annotation using EVidenceModeler and the Program to Assemble Spliced Alignments. Genome Biol 2008, 9:R7.

80. Spannagl M, Nussbaumer T, Bader KC, Martis MM, Seidel M, Kugler KG, Gundlach H, Mayer KF: PGSB PlantsDB: updates to the database framework for comparative plant genome research. Nucleic Acids Res 2016, 44:D1141–1147.

81. Ellinghaus D, Kurtz S, Willhoeft U: LTRharvest, an efficient and flexible software for de novo detection of LTR retrotransposons. BMC Bioinformatics 2008, 9:18.

82. Mistry J, Finn RD, Eddy SR, Bateman A, Punta M: Challenges in homology search: HMMER3 and convergent evolution of coiled-coil regions. Nucleic Acids Res 2013, 41:e121.

83. SanMiguel P, Gaut BS, Tikhonov A, Nakajima Y, Bennetzen JL: The paleontology of intergene retrotransposons of maize. Nat Genet 1998, 20:43–45.

84. Benson G: Tandem repeats finder: a program to analyze DNA sequences. Nucleic Acids Res 1999, 27:573–580.

85. Kurtz S, Narechania A, Stein JC, Ware D: A new method to compute K-mer frequencies and its application to annotate large repetitive plant genomes. BMC Genomics 2008, 9:517.

86. Wicker T, Yu Y, Haberer G, Mayer KF, Marri PR, Rounsley S, Chen M, Zuccolo A, Panaud O, Wing RA, Roffler S: DNA transposon activity is associated with increased mutation rates in genes of rice and other grasses. Nat Commun 2016, 7:12790.

87. Arend D, Junker A, Scholz U, Schüler D, Wylie J, Lange M: PGP repository: a plant phenomics and genomics data publication infrastructure. Database 2016, 2016.

88. Arend D, Lange M, Chen J, Colmsee C, Flemming S, Hecht D, Scholz U: e! DAL-a framework to store, share and publish research data. BMC bioinformatics 2014, 15:214.

89. Tange O: Gnu parallel-the command-line power tool. The USENIX Magazine 2011, 36:42–47.

90. Himmelbach A, Walde I, Mascher M, Stein N: Tethered Chromosome Conformation Capture Sequencing in Triticeae: A Valuable Tool for Genome Assembly. Bioprotocol 2018, 8:e2955.

91. S P, A H, M M, N S: In situ Hi-C for plants: an improved method to detect long-range chromatin interactions. In Plant long non-coding RNAs: methods and protocols. edited by J C, H-L W. New York, NY; 2019: Methods in molecular biology].

92. Mohamadi H, Khan H, Birol I: ntCard: a streaming algorithm for cardinality estimation in genomics data. Bioinformatics 2017, 33:1324–1330.

93. Himmelbach A, Ruban A, Walde I, Šimková H, Doležel J, Hastie A, Stein N, Mascher M: Discovery of multi-megabase polymorphic inversions by chromosome conformation capture sequencing in large-genome plant species. The Plant Journal 2018.

