## Supplementary Tables and Figures for "TRITEX: chromosome-scale sequence assembly of Triticeae genomes with open-source tools"

**Additional file 1**

**Supplementary Tables**

**Table S1.** Numbers of identified full-length retroelements in barley and wheat assemblies.

| TE family | Genome assembly | Solo-LTRs | Full-length | with TSD |
| --- | --- | --- | --- | --- |
| *RLC_Angela* | Chinese Spring DeNovoMagic | 6,935 | 21,948 | 19,379 (88%) |
| *RLC_Angela* | Chinese Spring TRITEX | 6,225 | 13746 | 12,177 (89%) |
| *RLG_Sabrina* | Chinese Spring DeNovoMagic | 656 | 604 | 492 (82%) |
| *RLG_Sabrina* | Chinese Spring TRITEX | 614 | 537 | 442 (83%) |
| *RLC_BARE1* | Morex V1 | 3,464 | 3,469 | 2,797 (81%) |
| *RLC_BARE1* | Morex V2 | 3,716 | 5,471 | 4,923 (90%) |

**Table S2: Classification of transposable elements in the Morex V1 and V2 assemblies.**

|  | Morex V1 | Morex V2 |
| --- | --- | --- |
| **Mobile Element (TXX)^1^** | **80.8** | **80.2** |
| **Class I: Retroelement (RXX)** | **75.2** | **74.2** |
| LTR Retrotransposon (RLX) | 75.0 | 74.0 |
| Ty1/copia (RLC) | 16.0 | 15.8 |
| Ty3/gypsy (RLG) | 21.3 | 21.3 |
| unclassified LTR (RLX) | 37.7 | 36.9 |
| non-LTR Retrotransposon (RXX) | 0.27 | 0.27 |
| LINE (RIX) | 0.25 | 0.26 |
| SINE (RSX) | 0.02 | 0.02 |
| **Class II: DNA Transposon (DXX)** | **5.28** | **5.64** |
| DNA Transposon Superfamily (DTX) | 5.03 | 5.39 |
| CACTA superfamily (DTC) | 4.74 | 5.11 |
| hAT superfamily (DTA) | 0.01 | 0.01 |
| Mutator superfamily (DTM) | 0.15 | 0.15 |
| Tc1/Mariner superfamily (DTT) | 0.02 | 0.02 |
| PIF/Harbinger (DTH) | 0.08 | 0.08 |
| unclassified (DTX) | 0.03 | 0.03 |
| DNA Transposon Derivative (DXX) | 0.20 | 0.20 |
| MITE (DXX) | 0.20 | 0.20 |
| Helitron (DHH) | 0.03 | 0.03 |
| unclassified DNA transposon (DXX) | 0.01 | 0.01 |
| Unclassified Element (TXX) | 0.32 | 0.31 |
| *Retro-TE/DNA-TE ratio* | *14.3* | *13.2* |
| *Gypsy/Copia ratio* | *1.3* | *1.4* |

^1^Except for the last two rows, numbers indicate the proportions (in percent) of the assembled genome sequence represented by the respective elements classes.

**Table S3: Intact LTR retrotransposons in the Morex V1 and V2 assemblies.**

|  | **Morex_V1** | **Morex V2** |
| --- | --- | --- |
| **Number of candidates** | 143,957 | 150,570 |
| **Number of high-confidence (HC) elements** | 31,643 (22.0 %) | 33,233 (22.1 %) |
| **RLG/ RLC ratio** | 0.87 | 0.91 |
| **Number of RLC elements** | 11,606 | 12,313 |
| **Number of RLG elements** | 10,116 | 11,210 |
| **Number of RLX elements** | 9,921 | 9,710 |

**Table S4: Numbers and sizes of sequence gaps in RLC_BARE1 elements in the Morex V1 and V2 assemblies.**

|  |  | **Morex v1** |  | **Morex V2** | | |
| --- | --- | --- | --- | --- | --- | --- |
| **Size range** | **Copies** | **Gaps per copy** | **Average gap length^1^** | **Copies** | **Gaps per copy** | **Average gap length** |
| 8000-8100 | 7 | 0.57 | 28 | 12 | 1.08 | 130 |
| 8100-8200 | 14 | 0.42 | 33 | 20 | 0.6 | 121 |
| 8200-8300 | 12 | 0 | 0 | 24 | 1.45 | 207 |
| 8300-8400 | 17 | 0 | 0 | 20 | 0.7 | 169 |
| 8400-8500 | 26 | 0.3 | 26 | 47 | 0.97 | 239 |
| 8500-8600 | 387 | 0.01 | 0 | 342 | 0.08 | 12 |
| 8600-8700 | 725 | 0.03 | 1 | 753 | 0.05 | 3 |
| 8700-8800 | 317 | 0.04 | 4 | 347 | 0.2 | 45 |
| 8800-8900 | 470 | 0.03 | 0 | 552 | 0.18 | 21 |
| 8900-9000 | 1353 | 0.02 | 2 | 792 | 0.17 | 23 |
| 9000-9100 | 31 | 0.87 | 124 | 225 | 1.19 | 240 |
| 9100-9200 | 16 | 0.87 | 56 | 268 | 1.24 | 259 |
| 9200-9300 | 22 | 1 | 105 | 265 | 1.29 | 299 |
| 9300-9400 | 20 | 1.14 | 130 | 340 | 1.29 | 307 |
| 9400-9500 | 13 | 1.23 | 39 | 329 | 1.22 | 351 |
| 9500-9600 | 19 | 1.1 | 186 | 235 | 1.51 | 478 |
| 9600-9700 | 12 | 0.75 | 120 | 157 | 1.71 | 528 |
| 9700-9800 | 8 | 0.62 | 135 | 160 | 1.83 | 622 |
| 9800-9900 | 0 | 0 | 0 | 152 | 2.09 | 710 |
| 9900-10000 | 0 | 0 | 0 | 151 | 2.21 | 760 |
| 10000-10100 | 0 | 0 | 0 | 155 | 2.3 | 859 |
| 10100-10200 | 0 | 0 | 0 | 124 | 2.19 | 835 |

^1^Average number of Ns per full-length copy in the respective size ranges

**Supplementary Figures**

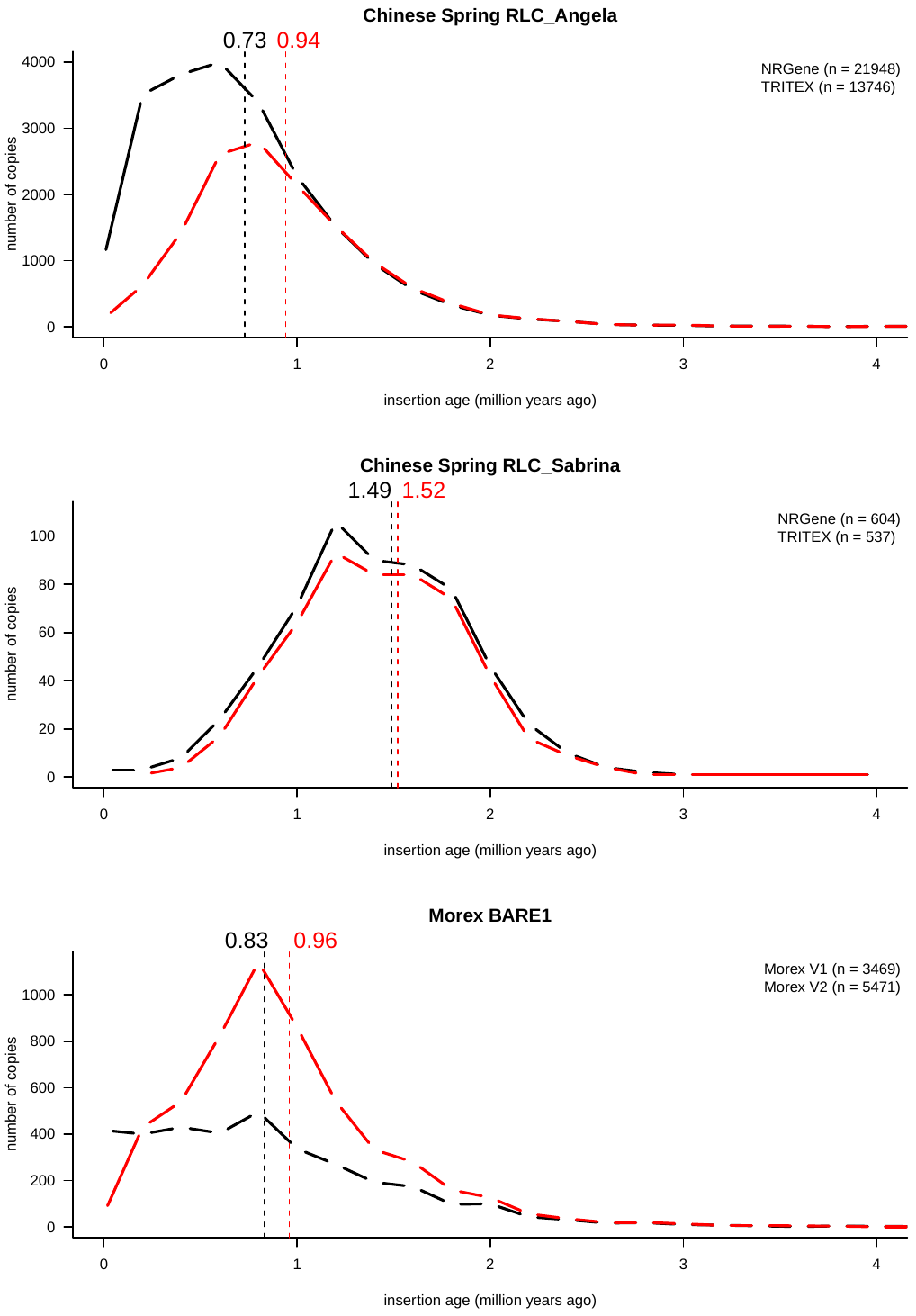

**Figure S1: Age distributions of full-length TEs in wheat and barley assemblies.** The medians of the distributions are indicated by dotted lines.

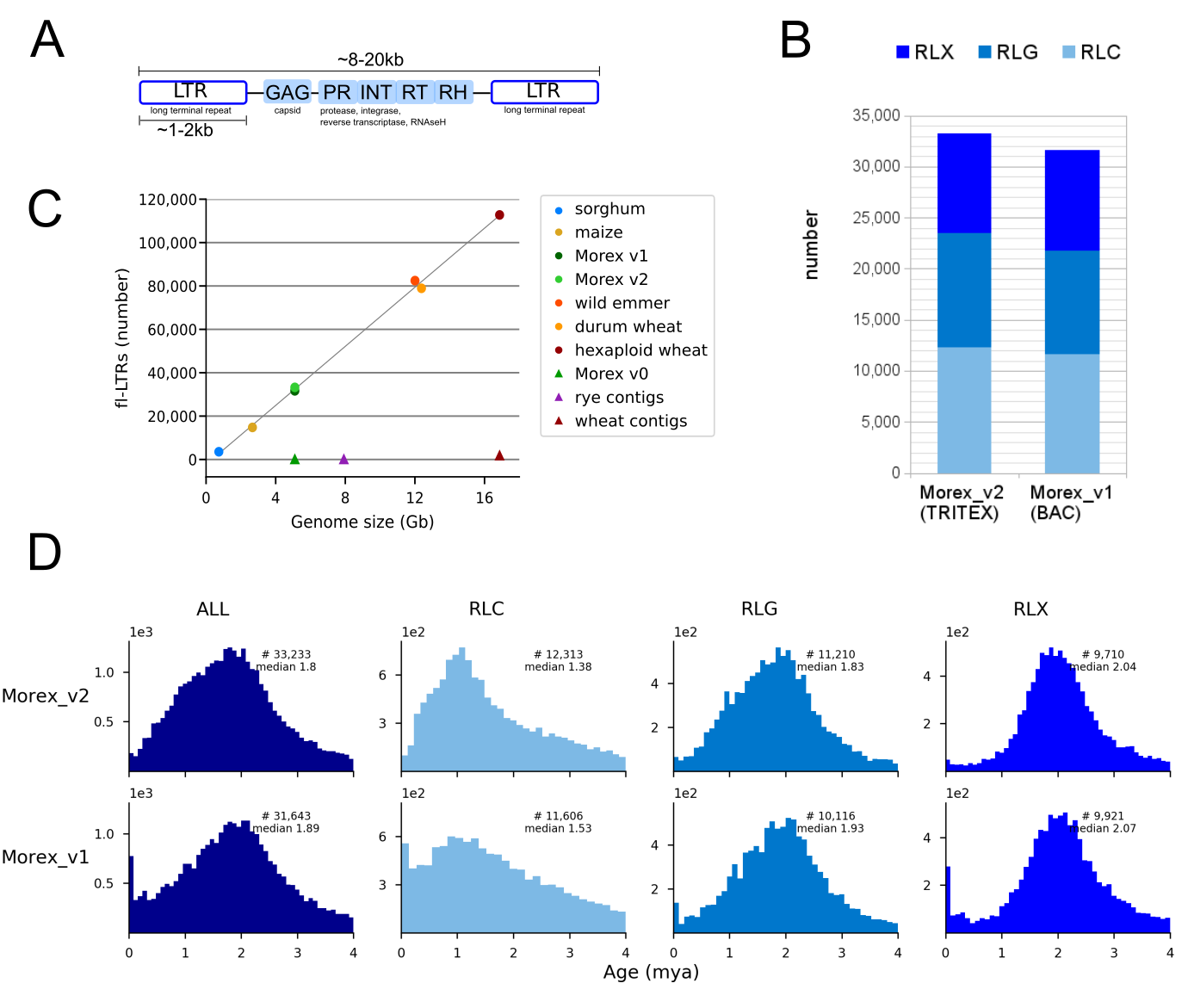

**Figure S2: Characterization of intact LTR-retrotransposons in the Morex V1 and V2 assemblies.** (A) Structure of LTR-retrotransposons. B) Numbers and superfamily distribution of quality filtered elements. RLC: Copia, RLG: Gypsy, RLX unassigned. (C) Linear relation between fl-LTR numbers and genome sized used as assembly quality metric. (D) Insertion age distribution of fl-LTRs: overall and per superfamily. The assemblies compared in panel C are the following: sorghum [1]; maize [2]; Morex v1, BAC-by-BAC assembly [3]; Morex v2 TRITEX assembly [this study]; wild emmer [4]; durum wheat [5]; hexaploid wheat, IWGSC RefSeqV1 [6]; Morex v0, fragmented WGS assembly [7]; rye contigs, WGS assembly [8]; wheat contigs, chromosome survey sequencing [9].

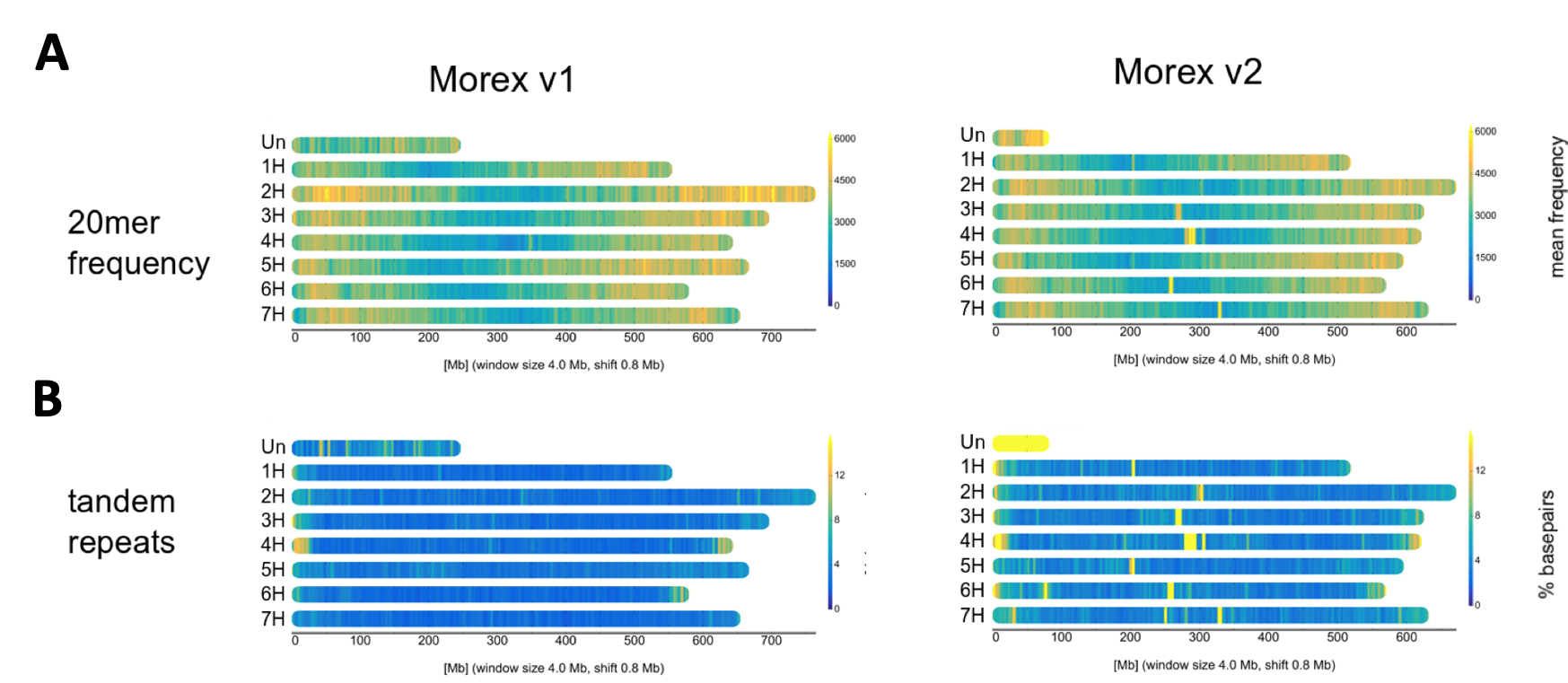

**Figure S3: Chromosome distribution of repeats in the Morex V1 and V2 assemblies.** (A) Frequencies 20-mer frequencies. (B) Locations of tandem repeats.

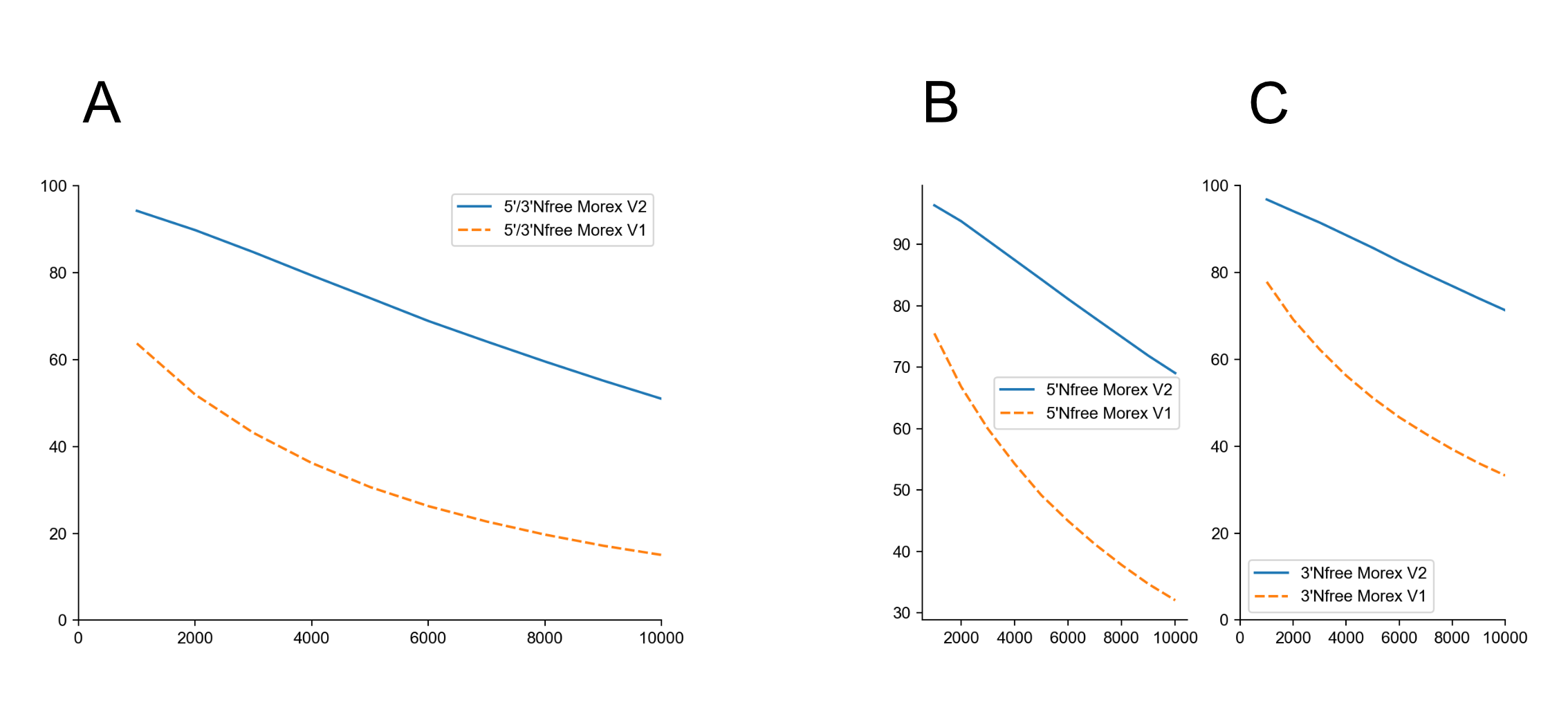

**Figure S4: Sequence gaps around genes in Morex V1 and V2 assembly.** The percentage of N-free flanking regions is plotted against the size of size of the flanking regions surrounding genes (A: combined 5’ and 3’ flanking regions; B:, 5’ flanking region; C: 3’ flanking region). The Morex V2 has longer N-free flanking regions than the Morex V1 assembly.

**Supplementary References**

1.	Paterson AH, Bowers JE, Bruggmann R, Dubchak I, Grimwood J, Gundlach H, Haberer G, Hellsten U, Mitros T, Poliakov A, et al: **The Sorghum bicolor genome and the diversification of grasses.** *Nature* 2009, **457:**551-556.

2.	Jiao Y, Peluso P, Shi J, Liang T, Stitzer MC, Wang B, Campbell MS, Stein JC, Wei X, Chin CS, et al: **Improved maize reference genome with single-molecule technologies.** *Nature* 2017, **546:**524-527.

3.	Mascher M, Gundlach H, Himmelbach A, Beier S, Twardziok SO, Wicker T, Radchuk V, Dockter C, Hedley PE, Russell J, et al: **A chromosome conformation capture ordered sequence of the barley genome.** *Nature* 2017, **544:**427-433.

4.	Avni R, Nave M, Barad O, Baruch K, Twardziok SO, Gundlach H, Hale I, Mascher M, Spannagl M, Wiebe K: **Wild emmer genome architecture and diversity elucidate wheat evolution and domestication.** *Science* 2017, **357:**93-97.

5.	Maccaferri M, Harris NS, Twardziok SO, Pasam RK, Gundlach H, Spannagl M, Ormanbekova D, Lux T, Prade VM, Milner SG, et al: **Durum wheat genome highlights past domestication signatures and future improvement targets.** *Nature Genetics* 2019.

6.	The International Wheat Genome Sequencing Consortium (IWGSC): **Shifting the limits in wheat research and breeding using a fully annotated reference genome.** *Science* 2018, **361:**eaar7191.

7.	International Barley Genome Sequencing Consortium: **A physical, genetic and functional sequence assembly of the barley genome.** *Nature* 2012, **491:**711.

8.	Bauer E, Schmutzer T, Barilar I, Mascher M, Gundlach H, Martis MM, Twardziok SO, Hackauf B, Gordillo A, Wilde P: **Towards a whole‐genome sequence for rye (Secale cereale L.).** *The Plant Journal* 2017, **89:**853-869.

9.	International Wheat Genome Sequencing C: **A chromosome-based draft sequence of the hexaploid bread wheat (Triticum aestivum) genome.** *Science* 2014, **345:**1251788.
